# Leveraging machine learning and accelerometry to classify animal behaviours with uncertainty

**DOI:** 10.1101/2024.12.28.630628

**Authors:** Medha Agarwal, Kasim Rafiq, Ronak Mehta, Briana Abrahms, Zaid Harchaoui

## Abstract

1. Animal-worn sensors have revolutionised the study of animal behaviour and ecology. Accelerometers, which measure changes in acceleration across planes of movement, are increasingly being used in conjunction with machine learning models to classify animal behaviours across taxa and research questions. However, the widespread adoption of these methods faces challenges from imbalanced training data, unquantified uncertainties in model outputs, shifts in model performance across contexts, and noisy classifications in continuous data streams, where predicted behaviours change abruptly within a sequence.
2. To address these challenges, we introduce an open-source approach for classifying animal behaviour from raw acceleration data. Our approach integrates machine learning and statistical inference techniques to evaluate and mitigate class imbalances, changes in model performance across ecological settings, and noisy classifications. Importantly, we extend predictions from single behaviour classifications to prediction sets: sets of behaviour labels guaranteed to contain the true behaviour with a pre-specified probability, in a framework analogous to the use of prediction intervals in statistical analyses.
3. We evaluate our approach via simulation and highlight its utility using data collected from a free-ranging large carnivore, African wild dogs (*Lycaon pictus)*, in the Okavango Delta, Botswana. We demonstrate significantly improved predictions along with associated uncertainty metrics in African wild dog behaviour classification, particularly for rare and ecologically important behaviours such as feeding, where correct classifications more than doubled following quality checks and data rebalancing introduced in our pipeline.
4. Our approach is applicable across taxa and represents a key step towards advancing the burgeoning use of machine learning to remotely observe around-the-clock behaviours of free-ranging animals. Future work could include the integration of multiple data streams, such as accelerometer, audio, and GPS data, for model training and could be incorporated directly into our pipeline.

## 1 Introduction

Animal-worn sensors have been pivotal in accelerating our understanding of animal ecology, providing a diversity of data that have advanced fundamental ecological theory and informed conservation actions (Snape et al., 2018; Nickel et al., 2021; VonBank et al., 2023; West et al., 2024). From their origins as tools primarily used to track animal locations and movements, animal-worn sensors have evolved to encompass a wide range of devices capable of monitoring the environments, behaviours, and internal states of animals (Wilmers et al., 2015). Animal-worn accelerometers – sensors that measure changes in acceleration across planes of movement – have been used to estimate energetic expenditure and infer animal behaviours for a wide variety of study systems and questions (see Halsey et al. (2011); Fehlmann et al. (2017)). From enabling the detection of spawning behaviours in large pelagic fish in the open ocean to characterising the hunting and energetics of elusive terrestrial predators (Wang et al., 2015; Clarke et al., 2021), accelerometers have become a valuable tool in ecology and have greatly expanded our capacity to understand ecological phenomena across spatial and temporal scales that were previously unattainable through direct observations alone (Studd et al., 2021).

The utility of accelerometers to capture animal behaviours lies in their ability to capture distinct waveform patterns within the raw data that correspond to specific movements or postures unique to different behaviours (Brown et al., 2013). Note, often the raw data are used rather than derived parameters, such as ODBA, due to the loss of potentially valuable information within the summary statistics. However, in contrast to the direct measurements provided by other sensor modalities, such as GPS or temperature sensors, the relatively abstract nature of accelerometer data can make the interpretation of waveforms challenging. As such, identifying behaviours with accelerometer data often requires pairing raw accelerometer data with known behaviours to create labelled datasets that can be used to learn the specific waveform patterns underlying different behaviours of interest (Brown et al., 2013). Due to the vast volumes of accelerometer data that can be collected and the subtle distinctions between behaviour signatures, manually detecting different behaviours within unseen accelerometer data can be challenging. To overcome this, machine-learning techniques are increasingly being leveraged to learn the accelerometer patterns underlying different behaviours from labelled datasets (Chakravarty et al., 2019; Garde et al., 2021; Otsuka et al., 2024), though ground-truthing remains essential.

Machine learning models for classifying animal behaviour encompass a diversity of techniques from classical machine learning classifiers, such as support vector machines (Martiskainen et al., 2009) and random forests (Lush et al., 2016), to more sophisticated models for sequential data, including convolutional neural networks, long short-term memory networks, and transformers (Otsuka et al., 2024). Yet current applications of these techniques in ecology have shared several challenges and limitations that can impact model performance and interpretability, including (1) imbalances in the volume of labelled data available for training models (i.e., class imbalance), (2) a lack of methods for statistically quantifying uncertainty in behaviour classifications, (3) shifts in model performance across application contexts (i.e., distribution shifts), and (4) rapidly fluctuating (and ecologically unlikely) classifications of consecutive behaviour segments. Below, we detail each of these challenges.

1. **Class imbalance**. Machine learning models for behaviour classification often require large volumes of labelled accelerometer data for each behaviour of interest for model training. Class imbalance, the unequal distribution of training data between the behaviours of interest, is pervasive in ecological datasets, where the frequency of common or easy-to-observe behaviours, such as resting or moving, can outweigh rarer behaviours, such as feeding or mating (e.g., Clermont et al. (2021); Otsuka et al. (2024)), and can bias model outputs (Johnson and Khoshgoftaar, 2019). However, existing approaches for addressing class imbalances typically aim for equal distributions across behaviour classes through oversampling the least represented behaviours (minority classes) or undersampling the most represented behaviours (majority classes), which can lead to biases in model outputs as a result of excessive resampling (Haixiang et al., 2017).
2. **Uncertainty quantification**. Most machine learning models provide single-label behaviour classifications without quantifying the uncertainties associated with the prediction (Nathan et al., 2012; Resheff et al., 2014). This lack of transparency limits our ability to assess a model’s limits and its applicability across datasets. Without clear uncertainty metrics indicating when and where model performance degrades, models can produce misleading results, particularly when applied to new datasets.
3. **Distribution shift**. Model performance can also decline due to distribution shifts, where the characteristics of the training dataset differ from the data that the models ultimately make their predictions upon (Kulinski and Inouye, 2023). For example, a model trained on one set of individuals within a population may perform poorly on future data from new individuals due to subtle behavioural differences, which could stem from morphological differences between animals in the population (Koh et al., 2021). Similarly, temporal changes, such as inter-annual behavioural variations, can degrade model performance when training and prediction periods differ (Ellington et al., 2020). Such distribution shifts are likely widespread within noisy ecological data and further underscore the importance of evaluation to detect and mitigate potential performance drops.
4. **Temporal context**. Finally, many (though not all) machine learning models classify behaviours based on isolated segments of accelerometer data, often ignoring the continuity or context of preceding behaviours (Riaboff et al., 2022). This lack of temporal context can lead to ecologically implausible behaviour sequences that rapidly fluctuate between distinct behaviours, undermining the biological realism of the classifications. Considering the accelerometry patterns before or after a specific segment of data can help improve prediction performance for that segment.

In this paper, we introduce an open-source approach for classifying animal behaviour using raw acceleration data that combines machine learning and statistical inference techniques to address these challenges. We start by introducing the concept of creating labelled training datasets and introducing a flexible rebalancing method that allows users to flexibly tune and evaluate the extent of resampling for their datasets. We then introduce a novel machine learning architecture for uncertainty quantification, wherein we train a prediction model (a convolutional neural network) and a conformal model for predicting animal behaviour with explicitly quantified uncertainties. Next, we describe how evaluation setups can be created to assess the presence of distributional shifts within datasets. Finally, we describe a technique for smoothing behaviour predictions across multiple temporal windows, leveraging smoothing techniques from signal processing and computer vision. We then apply our pipeline for predicting animal behaviour within a simulation study, where we provide a direct comparison of how model performance and outputs can improve with our contributions. Finally, we demonstrate the utility of our pipeline within real-world settings using accelerometer data collected from African wild dogs (*Lycaon pictus*) in the Okavango Delta, Botswana.

## 2 Classification Methods

This section outlines the machine learning methods used to classify animal behaviour from accelerometer data. We begin with the dataset construction process, detailing how raw sensor signals and behavioural annotations are processed to generate a labelled training dataset, including our approach to mitigating class imbalance. We then present the model architecture, including our approach to uncertainty quantification, and define the training loss function. Next, we describe how model performance is evaluated under various experimental setups to assess robustness to distribution shifts. Finally, we explain the smoothing of predictions to produce more biologically plausible behaviour sequences.

### 2.1 Data Preparation

#### 2.1.1 Creating Labelled Datasets

Acceleration data are collected as continuous streams of input from sensors mounted on animals, whereas behavioural annotations (i.e. labels of known behaviours with associated start and end timestamps) are determined by researchers based on video or other surveillance data (e.g., Fehlmann et al. (2017); Chakravarty et al. (2019); Clermont et al. (2021)). These disparate data sources are then carefully aligned temporally to create a labelled dataset of known behaviours and their associated acceleration readings from various animals and time intervals.

Given the timescales and sampling frequencies (which can exceed 100 Hz) of accelerometer deployments within ecology, the continuous stream of acceleration data, which can span from hours to years, is often too large to be loaded into CPU memory or even stored on disk as a single file. Therefore, in our pipeline, we first split continuous acceleration data into granular time segments of 12 hours each (i.e., half-days) to reduce CPU memory needs. Each half-day segment is accompanied by metadata, including details about the individual animal, observation day, and relevant environmental conditions. A comprehensive file is created, which stores the disk location of each segment along with its metadata. Our ML pipeline allows users to filter this metadata to retrieve segments that meet specific criteria. For instance, users can request segments from particular individuals for the training set and segments from another, disjoint set of individuals for the testing set in order to investigate whether the model can generalise to unseen individuals. This metadata-driven segmentation approach also supports the creation of train-test splits that reflect realistic distributional shifts between training and deployment, facilitating robust model evaluation (for detailed evaluation setups, see Section 2.4.2).

The duration of behaviours in a labelled dataset can vary significantly, whereas machine learning models are often trained using datasets with consistent time durations across examples. To address this, we extract windows of fixed duration from the labelled dataset. This fixed window duration, called window size, can be selected based on the time needed to detect accelerometer signals across the behaviours of interest and is usually a compromise between behaviours needing longer and shorter windows to capture their signals within accelerometer data. See Appendix B for details on selecting an appropriate window size and strategies for handling behaviours whose durations exceed or fall short of the chosen window size.

Given a sampling frequency of *f* Hertz and a window duration of *w* seconds, the resulting matrix for a tri-axial observation (across three spatial axes) has size *T* := ⌊*fw*⌋ (where ⌊*x*⌋ denotes the truncation of *x* to the nearest integer) along the temporal axis resulting in matrices of shape (3, *T*). The first dimension corresponds to the *x, y, z* acceleration axes. For *n* observations and *K* behaviour classes, we denote the labelled data by 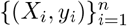, where *X*_*i*_ represents the windowed raw acceleration data with shape (3, *T*), and *y*_*i*_ is the behaviour label in the set {1, …, *K*} for *i*th observation. Herein, we will use the general terminology *input* to refer to each accelerometer clip *X*_*i*_ and *class* to each behaviour label *y*_*i*_. Once we have the labelled dataset 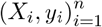, we randomly partition it into training, validation, and test datasets, maintaining the class distribution. However, we do not further stratify by attributes such as the individual from whom the signal was recorded or the specific recording time. Typically, we assume that these partitions follow a similar distribution, a condition commonly ensured in machine learning through random splitting.

#### 2.1.2 Class Rebalancing

With fixed-duration windows defined, we next address a common issue in behavioural datasets: class imbalance, which results in poor machine learning model performance on minority classes (Japkowicz and Stephen, 2002). We mitigate this issue by using flexible class rebalancing through resampling, adjusting the training data’s class distribution for improved class balance (Figure 1). Concretely, let *P*_*K*_ := (*p*_1_, …, *p*_*K*_) denote the empirical proportion of class labels, where *p*_*K*_ is the proportion of training examples that are of behaviour *k*. Ideally, the class distribution is the uniform distribution *U*_*K*_ := (1*/K*, …, 1*/K)*. We introduce the balancing parameter *θ* ∈ [0, 1] that controls the degree of resampling, creating the adjusted class distribution:

**Figure 1.**
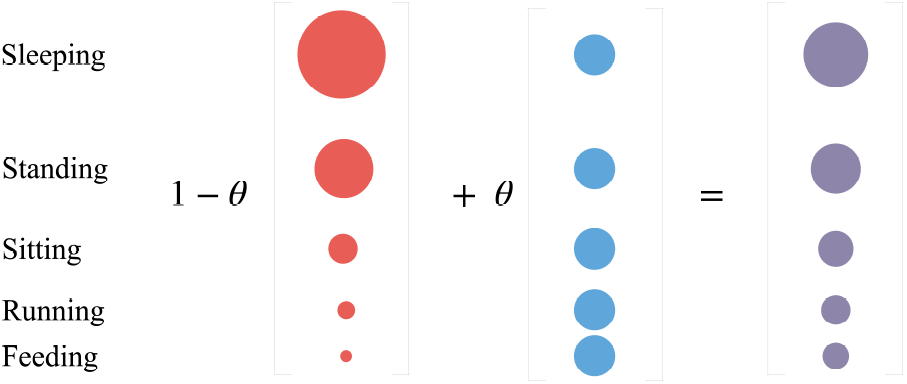
Rebalancing on a sample set of behaviour classes and their empirical distribution. The size of the red (left) dots represents the proportion of each behaviour class in the training dataset. The blue (middle) dots represent the ideal uniform class distribution (each class is equally represented). A linear combination of these two class distributions, parameterized by *θ*, gives a more balanced class distribution in violet (right).

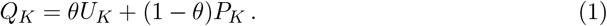

Here, *θ* = 0 retains the original distribution (no rebalancing), and *θ* = 1 corresponds to perfect class balance (identical sample sizes across classes). Higher *θ* values improve minority class representation but risk over-sampling the same observations. Conversely, lower *θ* values increase majority class representation, which may cause the model to favour the majority class and underperform on less frequent behaviours. For unbalanced datasets, we can select a non-zero *θ* and resample the training data to produce *Q*_*K*_, the adjusted class distribution. Once the adjusted class distribution *Q*_*K*_ = (*q*_1_, …, *q*_*K*_) is defined, we can draw *n*^′^ samples from the dataset with *Q*_*K*_ as the class distribution. To do this, we take *q*_*k*_ × *n*^′^ samples (with replacement) from the dataset for each behaviour class *k*, where *k* = 1, …, *K*. In our pipeline, users can specify their preferred *θ*, and this rebalancing is integrated into the training phase. The optimal *θ* should be fine-tuned for each dataset by training the model on rebalanced datasets across a range of equally spaced *θ* values and choosing the *θ* that gives the best average (across all behaviours) prediction accuracy on the validation set (see Figure 1).

### 2.2 Model Architecture and Uncertainty Quantification

With inputs preprocessed and rebalanced, we now describe the architecture of models used to predict behaviour and quantify uncertainty. Our approach combines two components: a *classification model* and a *conformal model*. Together, these models produce both a single most likely prediction and a prediction set that quantifies the uncertainty associated with that prediction.

The classification model, denoted by ℳ_1_ hereafter, takes a segment of accelerometer readings as input and outputs scores for each behaviour class. We call these scores *softmax scores* and they heuristically provide a probability distribution over the class labels (in that they are non-negative and add to one). A common practice is to choose the label with the highest softmax scores as our predicted label. This is called the *most-likely* prediction. However, the softmax scores are often miscalibrated (Guo et al., 2017), which means that they do not reflect the actual conditional probability of a label *y* being true, given the input *X*. In the absence of theoretical guarantees for the softmax scores, they can arbitrarily overfit or be completely random. For example, in an uncalibrated model, softmax scores of 0.9 versus 0.6 for the top label reflect relative model confidence but do not correspond to the true probability of class membership. Similarly, a lower score may simply indicate a more challenging or unfamiliar input, rather than a lower actual likelihood. The lack of formal probabilistic guarantees associated with the softmax scores output by ℳ_1_ is addressed via the conformal model. In multi-class classification, conformal prediction, denoted by ℳ_2_ hereafter, serves as a post-hoc calibration method that uses softmax scores from a base model (here ℳ_1_) to generate prediction sets: sets of class labels guaranteed to contain the true label with a pre-specified probability, typically denoted by 1 − *α*.

For the classification model, we employ an architecture called a one-dimensional convolutional neural network (1D CNN) to extract features from the raw accelerometry data for behaviour classification. For the conformal model, we use regularized adaptive prediction sets (RAPS) proposed by Angelopoulos et al. (2020). In the upcoming subsections, we describe technical details of the classification and conformal model architecture.

#### 2.2.1 Classification Model

We now describe the model architecture for ℳ_1_. Suppose *x* denotes the raw accelerometry input of size (3, *T*). The input signal is passed through *m* convolution layers. Each convolution layer consists of three steps: 1D convolution, max pooling, and activation. The input to a 1D convolution is of shape (*C*_in_, *T*), where *C*_in_ represents the number of input channels (*C*_in_ = 3 for tri-axial acceleration data), and the output is of shape (*C*_out_, *T*), where *C*_out_ represents the number of output channels. The 1D convolution is followed by a max pooling layer. Max pooling extracts the maximum value within a specified window of a feature map. Using a window size of 2, we halve the size of the feature map along the temporal axis. Finally, an activation layer is applied to the output of the max pooling layer. See technical details about convolution layers in Appendix A. We stack *m* convolution layers to obtain a sequentially deeper model (Figure 2). Standard LeNet-style architectures increase the number of output channels with convolution layers to increase model expressiveness. We double the number of output channels with each additional convolution layer, resulting in a final output of size (2^*m*−1^ *C*_out_, *T/*2^*m*^). This output is flattened and fed into a fully connected network to produce a vector **s** = (*s*_1_, …, *s*_*K*_) where recall *K* is the number of label classes. Finally, the softmax function converts this vector into class probabilities:

**Figure 2.**
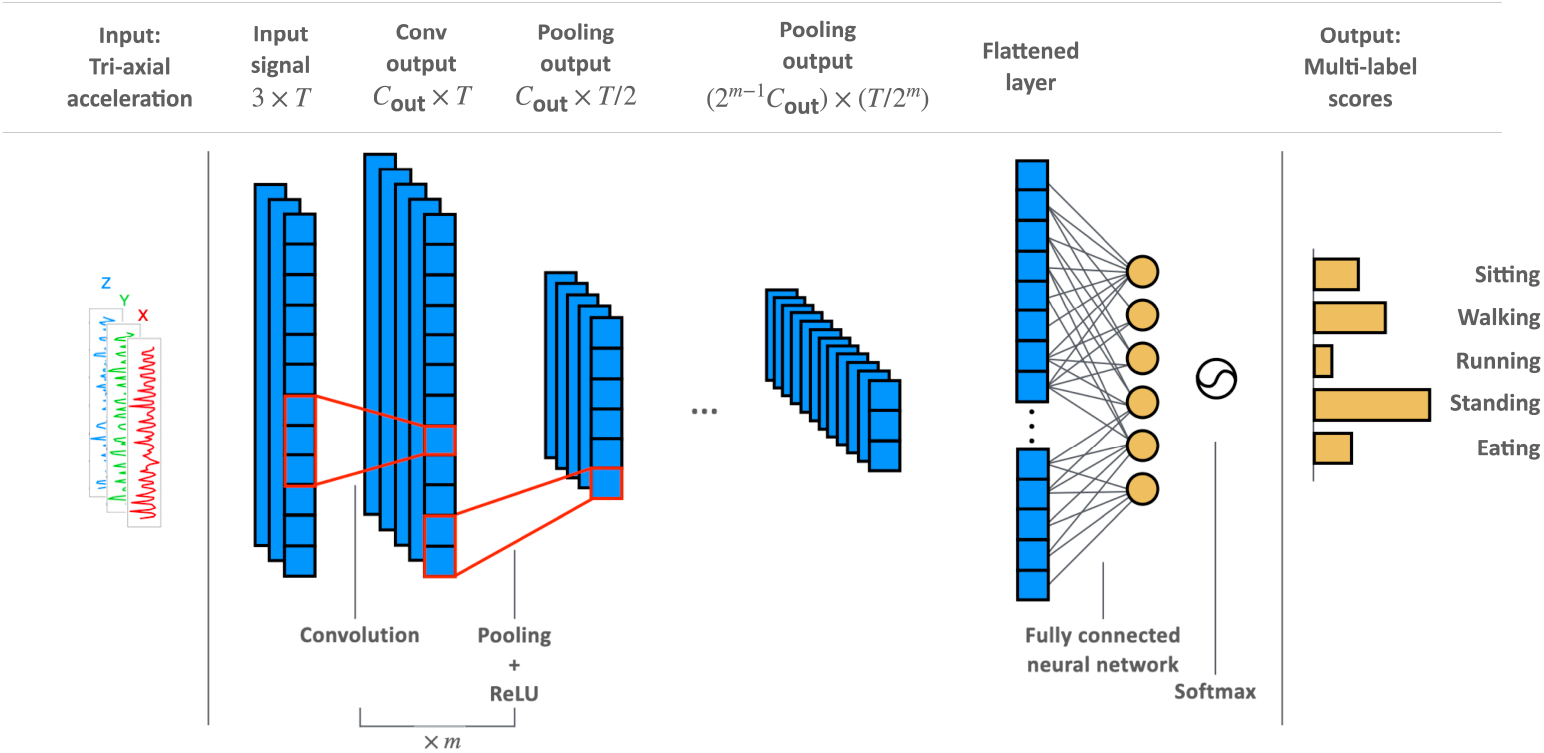
Schematic of a 1D convolution neural network. In the first convolutional layer, a 1D convolution is applied to an input of size (3, *T*), preserving the temporal dimension while increasing the number of channels to *C*_out_, a pooling layer reduces the temporal dimension by selecting maximum values within each window of size 2, followed by ReLU activation; applying ReLU(*u*) = max(0, *u*) elementwise. This process iterates in subsequent layers: **l**th convolution layer for **l* >* 1 receives a signal of size ((2^**l**−2^*C*_out_, *T/*2^**l**−1^)) and outputs features of size ((2^**l**−1^*C*_out_, *T/*2^**l**^)). The output of the last convolution layer is flattened into a 1D tensor, passed through a fully-connected network to yield a vector of size *K*. A softmax function (Eq (2) then converts this vector into a probability distribution, representing the probability of success for each class.

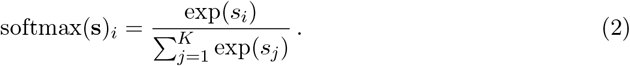

Thus, the final output of the 1D CNN is the softmax scores **s**, heuristically denoting the probability distribution over the class labels, as described in Section 2.2.

#### 2.2.2 Conformal Model

Constructing prediction sets requires two components: a trained base classifier and a held-out calibration dataset of moderate size, say *n*^′^ samples. We first define a prediction set and its coverage criterion mathematically. For a test observation (*X*_test_, *y*_test_), a set *C*(*X*_test_) is a valid prediction set with coverage (1 − *α*) ∈ [0, 1], say 90%, if the probability of *y*_test_ belonging to *C*(*X*_test_) satisfies the coverage property (Vovk et al., 2005; Vovk, 2012)

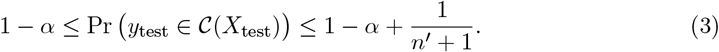

The function *C* is set-valued and is computed using the following steps: it takes an input *X*_test_, then uses the model ℳ_1_ (trained on the train set) to compute softmax scores, then calibrates these scores using the validation set (used as the calibration dataset), and then finally outputs a set of class labels based on the calibrated scores that satisfy the coverage criterion (3). Conformal model ℳ_2_ is responsible for the last two steps: i.e., calibrating the softmax scores output by ℳ_1_ and forming prediction sets (RAPS, satisfying (3)) from them. This procedure is detailed in Appendix C along with an exact algorithm for computing RAPS.

It is important to note that prediction sets are the sole output of conformal models; they do not provide probabilities for individual labels given the input signal *X*. Hence, they do not *correct* the softmax scores from ℳ_1_, instead, they calibrate and threshold them to produce prediction sets with guaranteed coverage. The coverage guarantee of RAPS, analogous to (3), is established in (Angelopoulos et al., 2023, Proposition 1). The entire calibration process, from computing calibrated softmax scores to constructing prediction sets, is encapsulated in the conformal model ℳ_2_.

### 2.3 Training Objective

Now we describe the *loss function* we minimise to train ℳ_1_. We minimise the popular one-vsall *binary cross-entropy loss*. Let *y*_*i,k*_ be the binary indicator for whether the *i*th observation belongs to class *k* and let *ŷ*_*i,k*_ denote the ℳ_1_’s output probability for sample *i* belonging to class *k*. The one-vs-all binary cross entropy loss for class *k* is

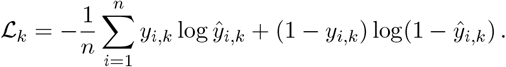

The overall loss is the sum of the terms above across all classes, i.e. we minimise

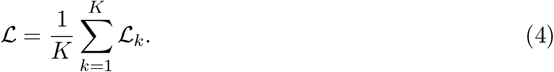

Minimising (4), ℳ_1_ produces softmax scores, which are plugged into ℳ_2_ as input to obtain RAPS.

### 2.4 Model Evaluation

In this section, we first outline the setup for assessing model robustness to potential distribution shifts in the dataset. Next, we define evaluation metrics for both most-likely predictions and prediction sets. Finally, we propose our approach for incorporating temporal context when evaluating our model on long-duration acceleration signals.

#### 2.4.1 Assessing Robustness to Distribution Shift

To assess the trained model’s performance, we then evaluate it on a subset of data not used for training ℳ_1_ and ℳ_2_, called the testing data. However, various temporal, environmental, intrinsic (such as body condition, sex, and age), and experimental factors can cause the future data that the model will encounter to differ slightly from the training data, leading to a distribution shift, which typically degrades model performance in real-world applications (Quiñonero-Candela et al., 2022). Therefore, it is critical to ascertain whether trained models are robust to distribution shifts using the available data. We achieve this by creating train-test splits that mimic expected distribution shifts by leveraging our pipeline’s ability to retrieve data subsets with specific properties using metadata (Section 2.1.1). For instance, to test robustness to individual-specific shifts, we might train on data from 4 out of 6 individuals and test on data from the remaining 2. Similarly, to evaluate robustness to temporal shifts, we can train on data collected before a specific year and test on data from subsequent years.

#### 2.4.2 Evaluation Metrics

After training the predictive and conformal models, ℳ_1_ and ℳ_2_, we proceed to evaluate their performance using specific metrics for each model. For ℳ_1_, we assess performance with three key metrics: precision, recall, and F1-score.

1. **Precision** measures how many of the examples the model labelled as a certain class actually belong to that class. Precision values range from 0 to 1, with 1 being perfect precision.
2. **Recall** measures how many examples belonging to a class were correctly labelled by the model, reflecting its ability to detect the presence of a class. Recall values range from 0 to 1, with 1 being perfect recall.
3. **F1-scores** aggregate precision and recall by taking their harmonic mean, given by the following formula

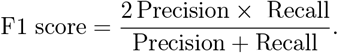

F1-scores offer a single metric that combines precision and recall to evaluate overall effectiveness. F1-score values range between 0 and 1, with 1 achieved only under perfect precision and recall. This metric is helpful in hyperparameter finetuning.

The evaluation metrics above are used to tune hyperparameters such as the number of CNN layers (*m*), the number of output channels *C*_out_, and the kernel size *h*. This tuning process involves training the model across a range of values for each hyperparameter and selecting the value that optimises the evaluation metric. To evaluate the conformal model ℳ_2_, which operates on top of the calibrated probability scores output by ℳ_1_, we use metrics such as average coverage and the average size of RAPS:

1. Coverage measures the proportion of instances for which the correct label is included in the prediction set provided by ℳ_2_. Coverage values range from 0 to 1, with 1 being perfect coverage.
2. Average RAPS size refers to the average size of prediction sets of the test data provided by ℳ_2_. Average RAPS size values range from 1 − *K*, where *K* is the total number of classes. Lower values of average RAPS size indicate more precise prediction sets.

#### 2.4.3 Smoothed Classifications for Temporal Context

After training and tuning the prediction model, we use the model to predict animal behaviours from a continuous stream of unseen accelerometry data. The model, trained on fixed window sizes *T*, breaks the acceleration data into chunks of width *T* and evaluates each chunk. Since the duration of each window is often chosen to be smaller than the true duration of behaviours, rapid transitions between classes can be biologically unlikely. To smooth these predictions, we average the predicted scores over a fixed number of windows, *s*, with a hop length *t*. Specifically, for an input signal *x* of shape (3, *N*), the signal is segmented into windows of width *T*, and the model produces a sequence of probability scores of shape (*K*, ⌊*N/T* ⌋). The windows can overlap to ensure that the behaviour scores do not vary too drastically between consecutive windows, and the degree of this overlap can be determined by the user.

To smooth sharp changes in these scores, we average *s* consecutive scores, calling this set the averaging window. Therefore, each averaging window averages scores over *s* × *w* seconds (recall *w* is the window duration in seconds). The averaging window shifts by *t* steps, causing consecutive averaged scores to share *s* − *t* evaluations. A larger *s* results in smoother, coarser behaviour predictions, suitable for longer activities like resting. Conversely, a smaller *s* provides finer, less smooth predictions, ideal for relatively shorter behaviours like eating, with the duration of the behaviour depending depending on the species and characteristics of the item being consumed (e.g., grazing on plants versus consuming a prey carcass). The value of *s* should be chosen based on the typical behaviour duration for the species studied. As an example, suppose we want to identify the occurrences of behaviour *k* in long sequences of acceleration data spanning months or even years. The typical duration of behaviour *k* in the concerned wildlife species is known to be *d*_*k*_, and the averaging window size should be a fraction of *d*_*k*_*/w*. Ideally, the averaged scores would favour behaviour *k* because it will receive the maximum vote for behaviour *k* from all *s* averaging windows. This approach ensures smoother and more reliable behaviour classification over long time horizons.

## 3 Simulation Results

In this section, we use a simulation study to demonstrate the application of our methodology across class imbalances, uncertainty quantification, distribution shifts, and temporal dependencies. Each observation is represented by the tuple (*X, Y*) ∈ *X* ×*Y*, where *X* denotes a tri-axial acceleration signal of shape (3, *T*): *T* = *fw* where *f* is the sampling frequency. Therefore, *X* ⊂ ℝ^3×*T*^. In these simulations, we consider *T* = 480. *Y* is the associated behaviour label. We consider four behavioural labels, namely *Y* = {feeding, moving, resting, vigilant}. Conditioned on a given behaviour *y* ∈ *Y*, the tri-axial acceleration signal is generated using a hierarchical model described in Appendix D.1. The specific hyperparameters for this model used to generate the train set are also detailed in Appendix D.1.

### 3.1 Class Rebalancing

In this simulation setup, we examine the impact of class rebalancing, as outlined in Section 2.1.2, by studying the dependence of model performance on the rebalancing parameter *θ* when the data-generating distribution has an extreme class imbalance. The data is generated with a highly imbalanced class distribution, where the proportion of the classes *feeding, moving, resting*, and *vigilant* are 2.5, 50.0, 45.0, and 2.5 percent, respectively. We sample 6000 (*X, Y*) pairs. Of these, 4000 samples form the train set, 1000 the validation set, and 1000 the test set. The CNN model is trained over a range of *θ* values to investigate the effect of rebalancing (Figure 3). All three evaluation metrics increased with *θ*, achieving the peak at *θ* values of 1.0, indicating this as the ideal amount of resampling that optimises model performance while minimising intensity of resampling (Figure 3). As a result, under *θ* values of 1.0, the prediction accuracy increased across all behavioural classes, increasing by more than four-fold for feeding, from 0.19 with no rebalancing to 0.85 with rebalancing.

**Figure 3.**
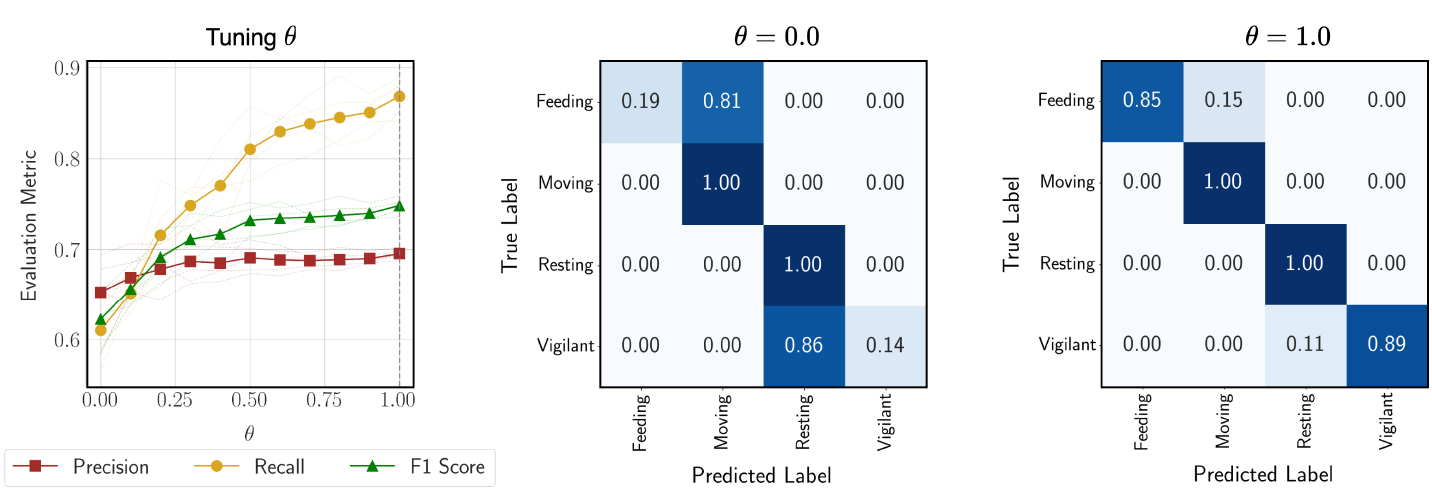
Left: Performance metrics (precision, recall, and F1 score) as a function of the rebalancing parameter *θ* on the validation set. Center: Confusion matrix for the test data when *θ* = 0.0 (no rebalancing). Right: Confusion matrix for the test data when *θ* = 1.0 (optimal rebalancing).

### 3.2 Uncertainty Quantification

In this subsection, we use the CNN trained with the optimal rebalancing parameter as our base prediction model ℳ_1_, and build a conformal model ℳ_2_ using the softmax outputs of ℳ_1_ evaluated on the validation set, as outlined in Section 2.2.2. We compared the accuracy of the most likely prediction, the coverage of RAPS, and the average size of RAPS among test samples for each behaviour (Table 1). Overall, 1.0% of all labels were incorrectly predicted using the single most likely prediction, with feeding being incorrectly predicted 15.4% of the time. However, upon fitting a conformal model with target coverage of 95% on the validation set, the true behaviour was identified within the prediction set on average 99.6% of the time (range: 92.3-100%), with feeding only being incorrectly predicted (not included in the prediction sets) 7.7% of the time. In fact, more than half of the behaviours (60%) that were incorrectly predicted using the single most-likely prediction had the true label present within the prediction set. Therefore, using prediction sets allowed us to improve the accuracy of our predictions by more than half. For example, a test observation with a true label feeding was incorrectly predicted as moving with a softmax score of 0.35. However, the prediction set for this observation was (Moving (0.35), Feeding (0.29)).

**Table 1:** Behaviour-wise summary of Top-1 prediction coverage, RAPS coverage, and average RAPS size.

| Behaviour | # Observations | Top-1 Coverage | RAPS Coverage | Avg. RAPS Size |
| --- | --- | --- | --- | --- |
| Feeding | 26 | 0.846 | 0.923 | 1.462 |
| Moving | 424 | 1.000 | 1.000 | 1.090 |
| Resting | 515 | 0.996 | 0.998 | 1.177 |
| Vigilant | 35 | 0.886 | 0.943 | 1.257 |

### 3.3 Assessing Robustness to Distribution Shift

In this simulation setup, we aimed to quantify how a trained CNN’s predictive performance degrades gracefully as the distribution shift increases. While keeping the distribution of the train set fixed (Appendix D.1, we parametrise the shift between the train and test set using a single scalar parameter *ε >* 0 in the manner described in Appendix D.2. At *ε* = 0, the train and test distributions are identical. As *ε* increases, the test distribution diverges due to changes in the conditional distribution of *X* given *Y*, particularly making behaviour pairs feeding-moving and resting-vigilant increasingly indistinguishable.

To quantify this divergence, we compute the sliced Wasserstein distance (SWD) (Bonet et al., 2022) between empirical train and test distributions. However, high dimensionality poses challenges: with *T* = 480, each *X* ∈ ℝ^3×*T*^ yields a 1440-dimensional vector after flattening, which is unsuitable for reliable SWD estimation. To mitigate this, we extract 34 summary features per observation based on established ecological methods (Martiskainen et al., 2009; Nathan et al., 2012), resulting in dimension-reduced train and test datasets. Full details on feature construction and SWD computation are provided in Appendix D.2.

We train ℳ_1_ on 5000 train samples with a rebalancing parameter of *θ* = 0.8. For model evaluations, we sample 5000 samples from the test distribution for a range of values of *ε*. Let the SWD between the *X* marginal of train distribution (*P*_*X*_) and test distribution (*Q*_*X*_) be denoted by *D* (*P*_*X*_, *Q*_*X*_). As expected, increasing *ε* induces a growing shift between *P*_*X*_ and *Q*_*X*_, evidenced by rising *D* (*P*_*X*_, *Q*_*X*_) in Figure 4 (left). A CNN trained on the training data exhibits progressively poorer performance as *ε* increases. This degradation is reflected in the declining precision, recall, and F1 scores shown in the centre and right panels of Figure 4. Confusion matrices (Figure 5) for *ε* = 0.1, 0.2, and 0.5 further underscore the performance drop due to distribution shift with increasing *D* (*P*_*X*_, *Q*_*X*_). Thus, our method provides a reliable means of identifying problematic distribution shifts.

**Figure 4.**
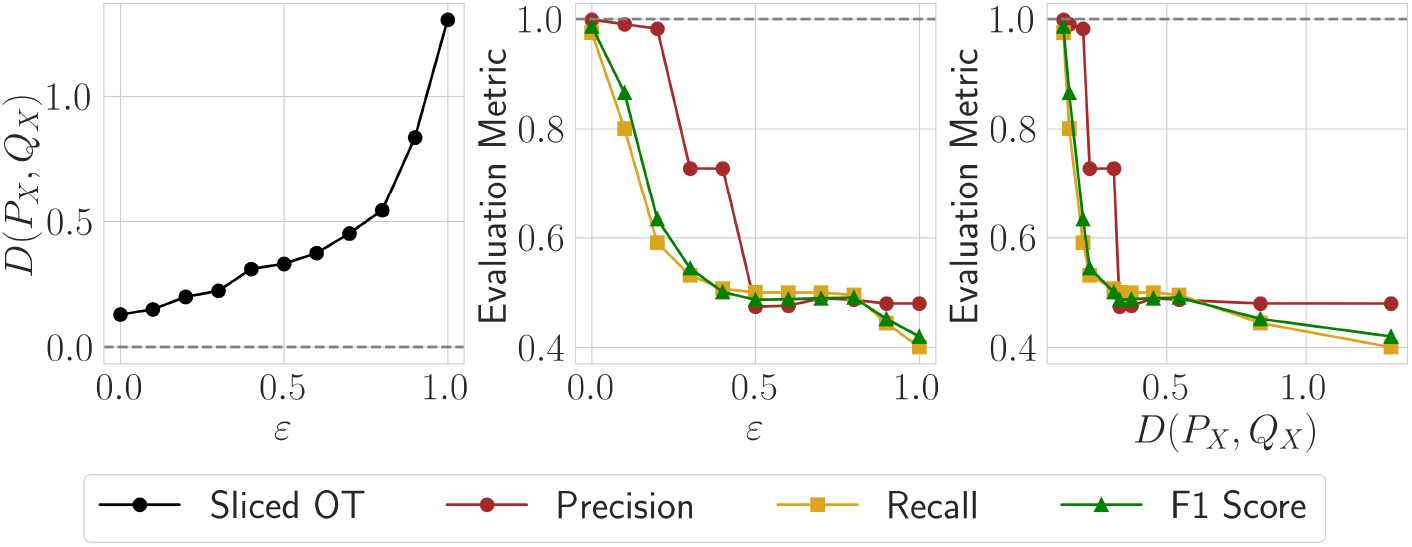
Left: Sliced Wasserstein distance (*D*(*P*_*X*_, *Q*_*X*_)) is a measure of discrepancy between the *X* marginal of train data (denoted by *P*_*X*_) and test data (denoted by *Q*_*X*_). Center: The decay of model performance metrics - precision, recall, and F1 score as *ε* increases. Right: The decay of model performance metrics as *D* (*P*_*X*_, *Q*_*X*_) increases.

**Figure 5.**
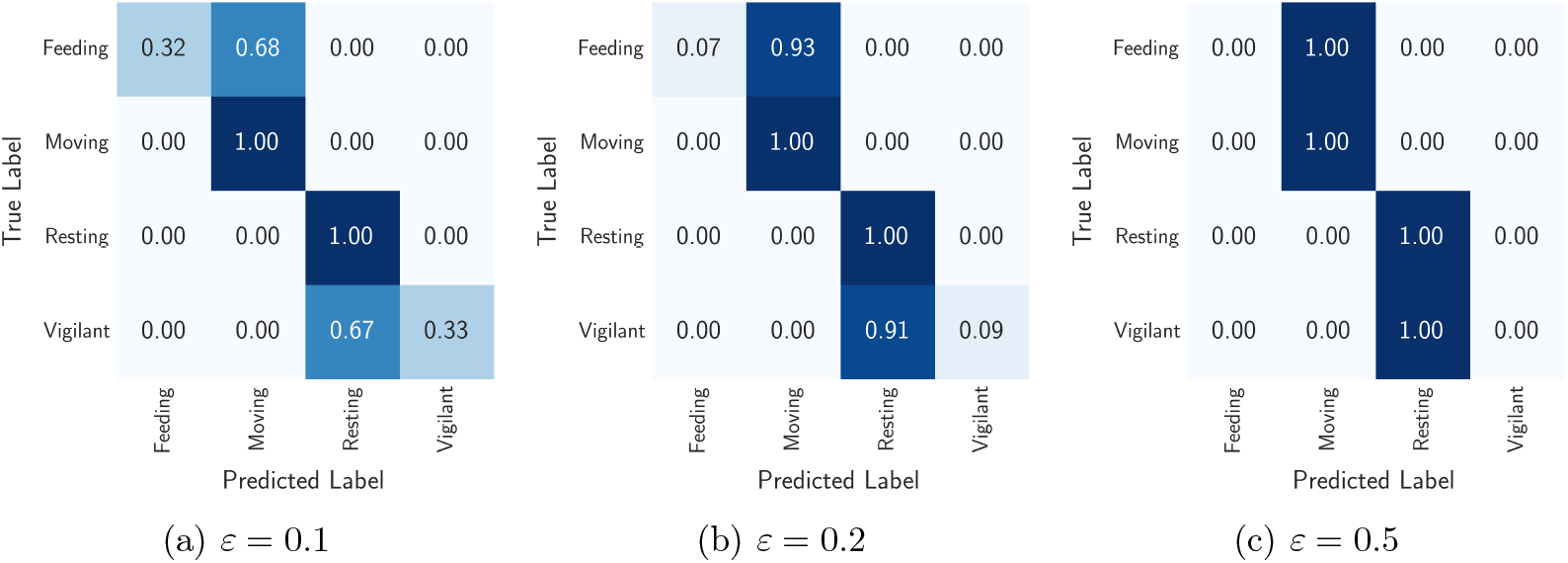
Confusion matrices for the trained CNN’s performance on the test data with distributed shift quantified by energy score of 0.150 (*ε* = 0.1), 0.200 (*ε* = 0.2), and 0.331 (*ε* = 0.5).

### 3.4 Smoothed Classifications for Temporal Context

To demonstrate the effect of temporal smoothing, outlined in Section 2.4.3, we simulated a 24-hour sequence of acceleration and behaviour data using a Markov transition model (see Appendix D.3) and introduced snippets of corrupted signals, which may reflect sensor malfunctions during real-world data collection. We introduced artificial corruptions, but with distinct generation hyperparameters from true signals. The duration of each corrupted signal snippet is randomly sampled between 2*w* and 4*w*. Figure 6 (top) displays an exemplary corrupted signal. The simulated signal was segmented into non-overlapping *w*-duration windows, and per-segment softmax scores were computed. These raw scores (Figure 6, middle) reveal that model ℳ_1_ erroneously assigns the *moving* behaviour to corrupted windows, causing score spikes for this class. To mitigate these errors, scores were temporally smoothed by averaging over *s* = 20 consecutive window (with *w* = 30 seconds, this is averaging over 10 minutes). The smoothed scores are shown in Figure 6 (bottom).

**Figure 6.**
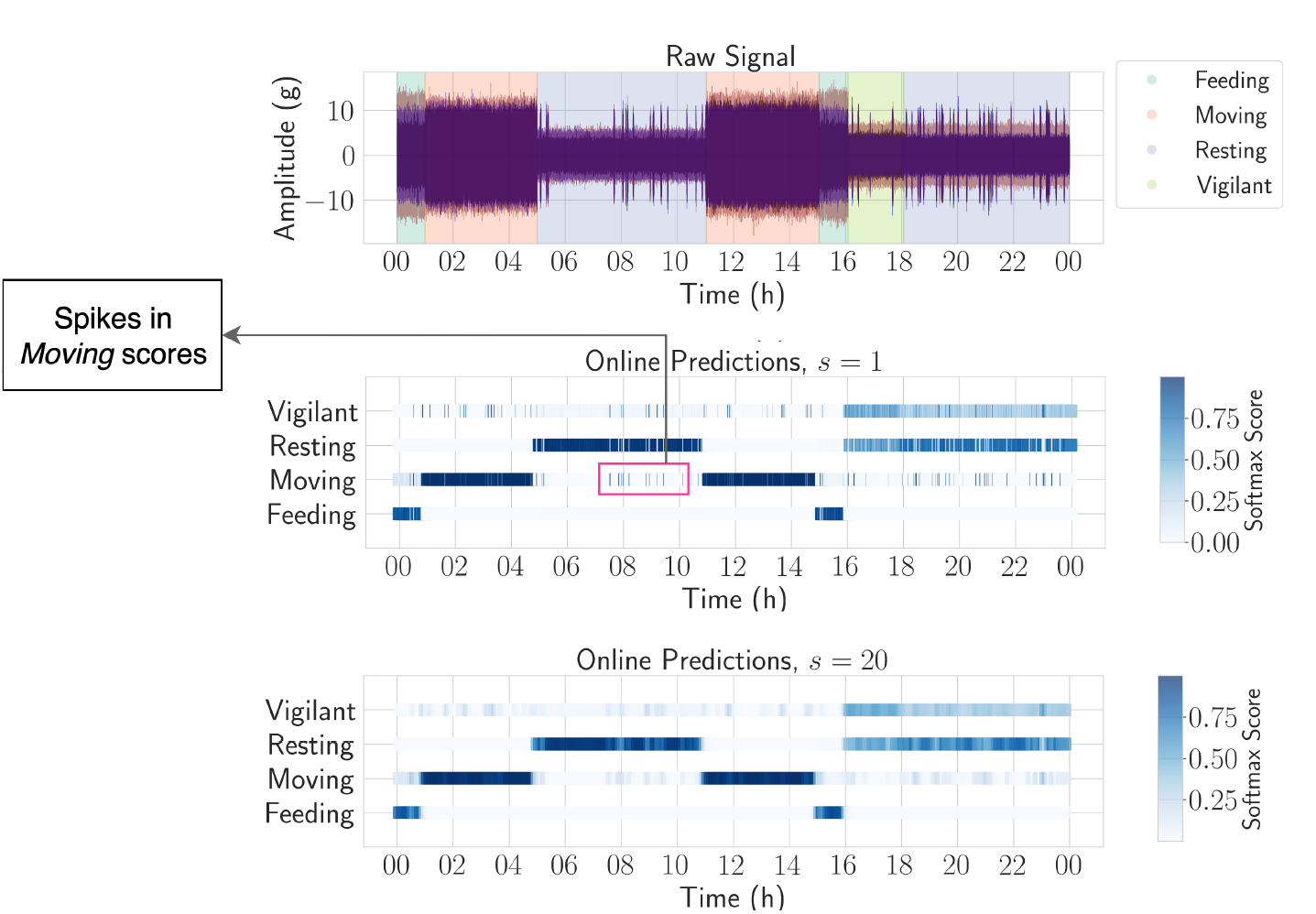
Top: Simulated raw tri-axial acceleration signals (X, Y, Z) with behavioural annotations over a 24-hour period. Middle: Softmax scores predicted by model ℳ_1_ for each 30-second window, plotted along the temporal axis. Bottom: Smoothed softmax scores obtained by averaging over 20 consecutive windows (10 minutes).

Ideally, softmax scores should remain stable within a single behaviour episode, particularly for behaviours that the model predicts with high confidence. For behaviours that the model tended to confuse, such as feeding and moving, we expected higher variability in their respective softmax scores, while scores for resting should remain consistently low. Beyond this expected pattern, we observed unexpectedly high variability in the *moving* scores not only during true *moving* or *feeding* episodes (which the model often confuses with *moving*) but also during *resting* and *vigilant* periods. This suggests that corrupted signals inflate the variability of *moving* scores across all behaviours.

Quantitatively, the relative standard deviation (SD) of moving scores, defined as the ratio of the SD of moving scores when another behaviour is true to the SD when moving is the true behaviour, was 1.46, 1.60, and 1.29 for feeding, resting, and vigilant, respectively. After applying smoothing, the corresponding values were 1.98, 0.84, and 0.40, reflecting reduced variability when the model does not confuse between a behaviour and moving (resting and vigilant). Ultimately, without smoothing, corrupted signal segments were often misclassified as moving, leading to erratic spikes during periods of true resting behaviour (Figure 6). These fluctuations were substantially reduced with smoothing. Notably, the variability of moving still showed elevated variance during feeding episodes due to the model’s inherent confusion between feeding and moving, but smoothing effectively mitigated the influence of the corrupted signal on the variability of moving scores during other behaviour episodes. An extended table reporting the standard deviations of softmax scores for each behaviour and its discussion is provided in Appendix D.4. Thus, temporal smoothing allowed us to use context from surrounding windows to reduce the impacts of corrupted signals and noisy classifications.

## 4 African Wild Dogs Study

In this section, we apply our method to a case study on African wild dogs. We describe the data collection process, including collar deployments, audio and video recording, and behavioural annotation. We explain how these data were processed to create a labelled training set and discuss context-specific challenges, such as class imbalance and behavioural ambiguity. Throughout, we highlight where the general pipeline described in Section 2 was adapted or applied to this ecological system.

### 4.1 Data Collection and Annotation

From October 2021 to September 2023, we deployed wildlife tracking collars with inbuilt GPS and accelerometer sensors (GPS PLUS 1C, Vectronic Aerospace, Germany) on five free-ranging African wild dogs (*Lycaon pictus*) in the Okavango Delta, Botswana (centre 19^◦^31*S*, 23^◦^37*E*). The study area encompassed approximately 2, 600km^2^ and was a mosaic of habitat types ranging from seasonally flooded floodplains to mixed woodlands, with little variation in elevation (for more details, see Rafiq et al. (2023b)). All capture and collaring was approved by the University of Washington Animal Care and Use Committee (Protocol #4514-01) and the Botswana Department of Wildlife and National Parks.

Tracking collars were scheduled to record GPS fixes every three hours and continuous accelerometer data at 16 Hz. Animal-worn microphones (uMoth, Open Acoustic Devices, United Kingdom) were attached to collars with a bespoke attachment mechanism that detached from collars following battery expiry after three weeks (Rafiq et al., 2023a). Microphones were included in collar deployments as the recordings could be used to collect labelled data of behaviours for the accelerometers. To maximise deployment durations, microphones were scheduled to record audio at 16 kHz in 25-second on and 5-second off sampling cycles between 0600-1000 and 1600-2000 (local time). Recording windows were chosen to correspond with the times that African wild dogs were most active (Rafiq et al., 2023b). Further details on collar deployments can be found within the supplemental material.

We labelled 72.83 hours of the acceleration data into one of five biologically relevant behavioural classes – *feeding, resting, moving, running*, and *vigilant* – by first pairing data with timestamped behaviours observed within our vehicle-collected video (see Table 8 in the appendix for a description of each behaviour). Herein, we refer to our five behavioural class labels from the video as *video labels*. We selected these behavioural classes based on their relevance to wider ecological questions of interest and as they represented distinct behaviours possible to unambiguously identify within the acceleration signals, based on posture and movement. Behaviours that did not fit into these categories or that were unclear from video recordings were excluded, and in total, we discarded a total of 1.28 hours of labelled data (1.23 % of our collected labels) spanning ten additional behaviours (e.g., digging, drinking, scent marking).

There was significant class imbalance within our behaviours due to the relative rarity of less observed behaviours, as well as the logistical challenges of directly observing these behaviours in the wild from a vehicle. Consequently, after using the video-labelled audio data to listen to the audio profiles of our behaviours, we used a subset of our audio (for which we had no video) to add additional feeding, moving, running, and vigilance labels and matched them to our acceleration data. We used audio to only add labels for these behaviours because these could be clearly delineated within the audio datasets with high levels of certainty. Herein, we refer to these as *audio labels*. In total, we collected 491.07 hours of audio data, only a small portion of which we listened to for labelling (5% of total audio collected) due to data processing times. At the end of this step, we had 73.94 hours of labelled acceleration data (both video and audio labels) with labels corresponding to one of the five chosen behavioural classes. Instances of this labelled acceleration data, of variable duration, are referred to as “matched segments”.

We further discarded matched segments under one-second in duration because they were too brief to capture meaningful acceleration patterns, and we removed matched segments of feeding, moving, and running under eight seconds after visual inspection of waveforms suggested such labels were unreliable. This amounted to removing approximately 0.02% of the filtered data and left us with 72.21 hours of labelled accelerometer data for our five behaviours (Table 2), with the most represented behaviour (resting) having ≈ 90 times more duration of matched accelerometry signal than the least represented behaviour (running). We used this filtered dataset as our training data. (Table 2). The matched behaviour-acceleration pairs exhibited significant variability in duration due to inherent differences in the durations of behaviours exhibited by animals and the availability of reliable video recordings (see Table 9 in Appendix E for an individual-wise data summary).

**Table 2:**
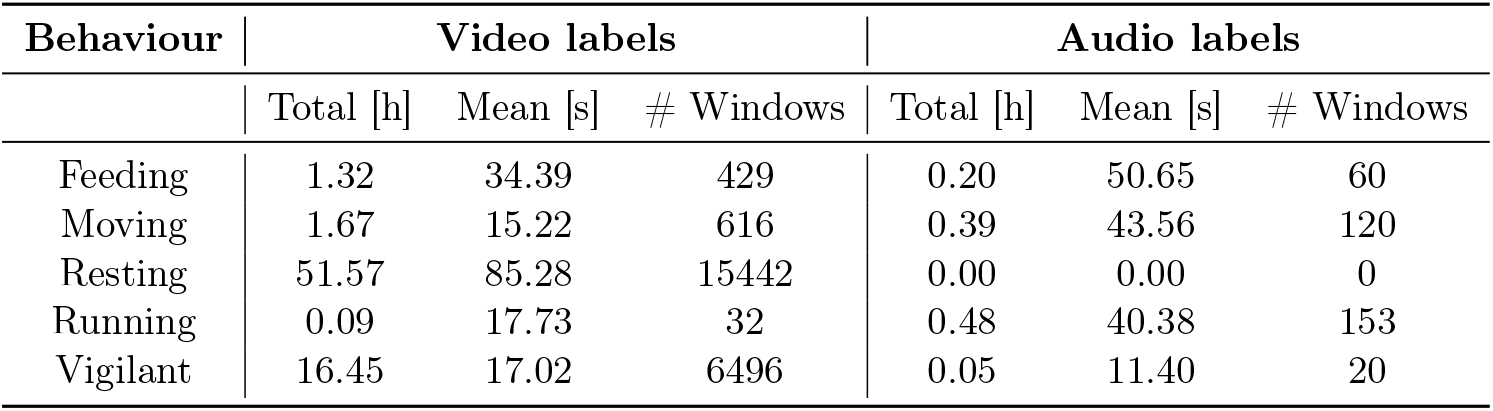
Behaviour-wise summary of the filtered 72.21 hours of labelled acceleration data. The table shows the total duration in hours (column name: Total [h]), mean duration in seconds (column name: Mean [s]), and number of extracted fixed-duration windows (column name: # Windows) for each behaviour across the video and audio labels.

| Behaviour | Video labels |  |  | Audio labels |  |  |
| --- | --- | --- | --- | --- | --- | --- |
|  | Total [h] | Mean [s] | # Windows | Total [h] | Mean [s] | # Windows |
| Feeding | 1.32 | 34.39 | 429 | 0.20 | 50.65 | 60 |
| Moving | 1.67 | 15.22 | 616 | 0.39 | 43.56 | 120 |
| Resting | 51.57 | 85.28 | 15442 | 0.00 | 0.00 | 0 |
| Running | 0.09 | 17.73 | 32 | 0.48 | 40.38 | 153 |
| Vigilant | 16.45 | 17.02 | 6496 | 0.05 | 11.40 | 20 |

### 4.2 Training Data

The duration of labelled behaviours varied significantly within and across behaviour classes in the filtered dataset. For instance, among the video labels, the average duration of resting was 85.28 seconds, whereas the average duration of running was 17.73 seconds. We extracted fixed-duration “windows” from matched behaviour-accelerometer pairs, following the methods described in Section 2.1.1. We determined the window length as the median duration of all behaviours, resulting in *w* = 12.94 seconds, following sensitivity analyses where we ran our pipeline for different percentiles of window duration, 10th to 90th, in 10-percentile increments. We selected the median duration of all behaviours as our window size based on a balance between model performance and computation times. Shorter window durations, while increasing the number of observations, also required more extensive data segmentation and longer model training times. Illustrating this balance, a shorter window based on the 30th percentile (6.93 seconds) yielded comparable model performance to our chosen 12.94-second window but increased overall computation times threefold. Our chosen *w* = 12.94 resulted in 23, 368 behaviour-acceleration pairs (Table 2) (see Figure 11 in Appendix E for representative waveforms for each behaviour class). The behaviours feeding, moving, resting, running, and vigilant accounted for approximately 2.09, 3.15, 66.08, 0.79, and 27.88 percent of the dataset, making the class distribution unbalanced.

For training the 1D CNN for behaviour classification, we divided our 23, 368 matched acceleration behaviour pairs into train and test sets. By default, we split the dataset into train and test sets using an 80-20% ratio, preserving the class distribution across both. The train set was further divided into train and validation subsets (also 80–20%), maintaining class balance. The train data was used to train ℳ_1_, while the validation data was used as the calibration set to fit ℳ_2_. We also explored three alternative train-test splits, each defined by specific criteria applied to the training and test samples. These splits did not yield fixed train–test size ratios, as the number of observations meeting the respective criteria varied. For instance, in an individual-based split, where the test set consisted of data from a single individual and the training set comprised data from all others, the resulting train-test ratio was approximately 70-30%. Further details on these distribution shifts and the corresponding train–test splits can be found in Section 4.5.

### 4.3 Class Rebalancing

Our dataset had a high class imbalance, with minority classes - feeding, moving, and running - comprising only 6% of the data. As described in Section 2.1.2, we rebalanced the training data using *θ* values from 0.0 to 1.0 (Figure 7) and evaluated performance using precision, recall, and F1 scores on the validation set. F1 score increased with *θ*, achieving the maxima at *θ* values of 0.3. Thus, the *θ* value was tuned to 0.3, representing the ideal amount of resampling for optimising model performance while minimising the extent of resampling adjustments. The resultant class distribution obtained by resampling with *θ* = 0.3 improves the proportion of behaviours feeding, moving, resting, running, and vigilant to approximately 7.47, 8.16, 52.46, 6.53, and 25.38 percent in the train set. The confusion matrices in Figure 7 show how class rebalancing successfully uplifts the prediction performance of the model, especially for the minority class feeding. Specifically, following rebalancing, the proportion of correctly predicted feeding instances in the test set (true positives) increased from 0.62 to 0.97.

**Figure 7.**
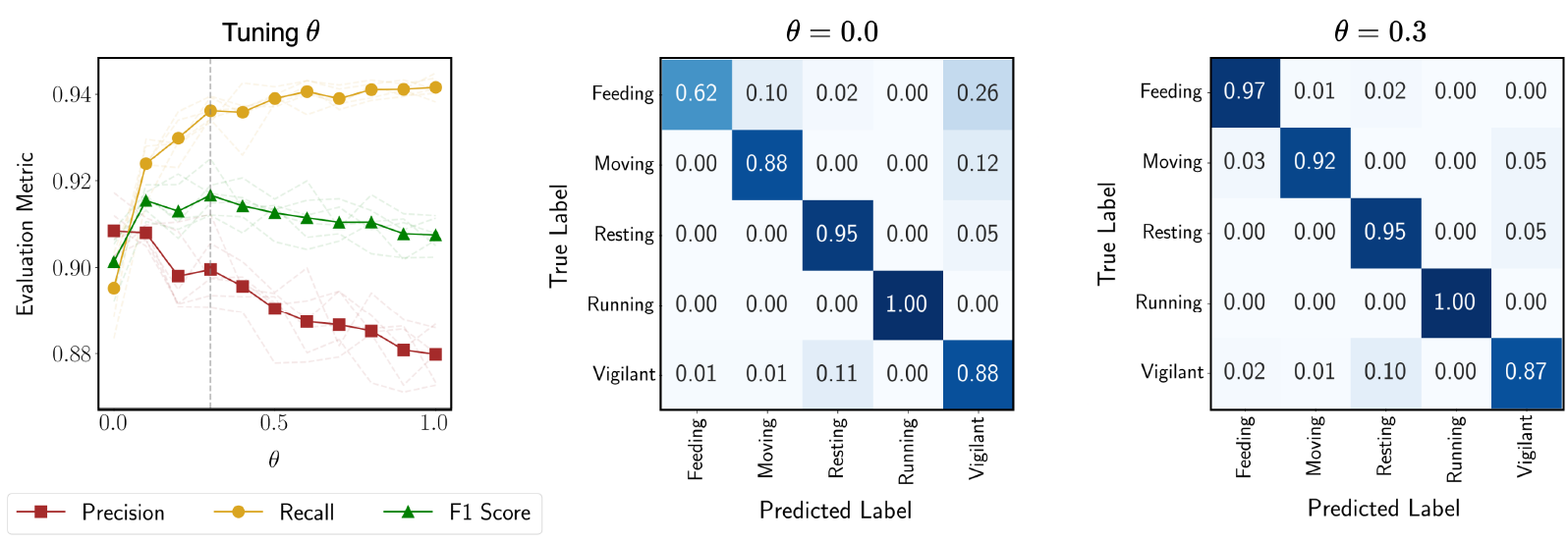
Summary showing how flexible class rebalancing improved model performance when the original training data comprises only 2.09% of Feeding samples. The left plot illustrates how precision, recall, and the F1 score (averaged over training runs with six different seeds) evolved with increasing rebalancing intensity (parameterized by *θ*) when the trained model is evaluated on the validation set. The confusion matrices compare classification results on the test set: the first matrix for no rebalancing (*θ* = 0.0) and the second for optimal rebalancing (*θ* = 0.3).

### 4.4 Model Training

After class rebalancing, we trained the prediction model ℳ_1_ using our 1D CNN architecture (see Section 2.2), minimising the multi-label loss Eq (4). The hyperparameters of the model were tuned by comparing model performance for different combinations of hyperparameter values on a held-out validation set. Refer to Appendix F.1 for details and recommendations on hyperparameter fine-tuning. The class distribution in the validation was maintained to be the same as that of the remaining training set. We fit the conformal model ℳ_2_ on the output probabilities of the prediction model with confidence level 1 − *α* = 0.95. Figure 8 illustrates the RAPS for a random test sample for each behaviour class, along with the associated probability of each class in the prediction set.

**Figure 8.**
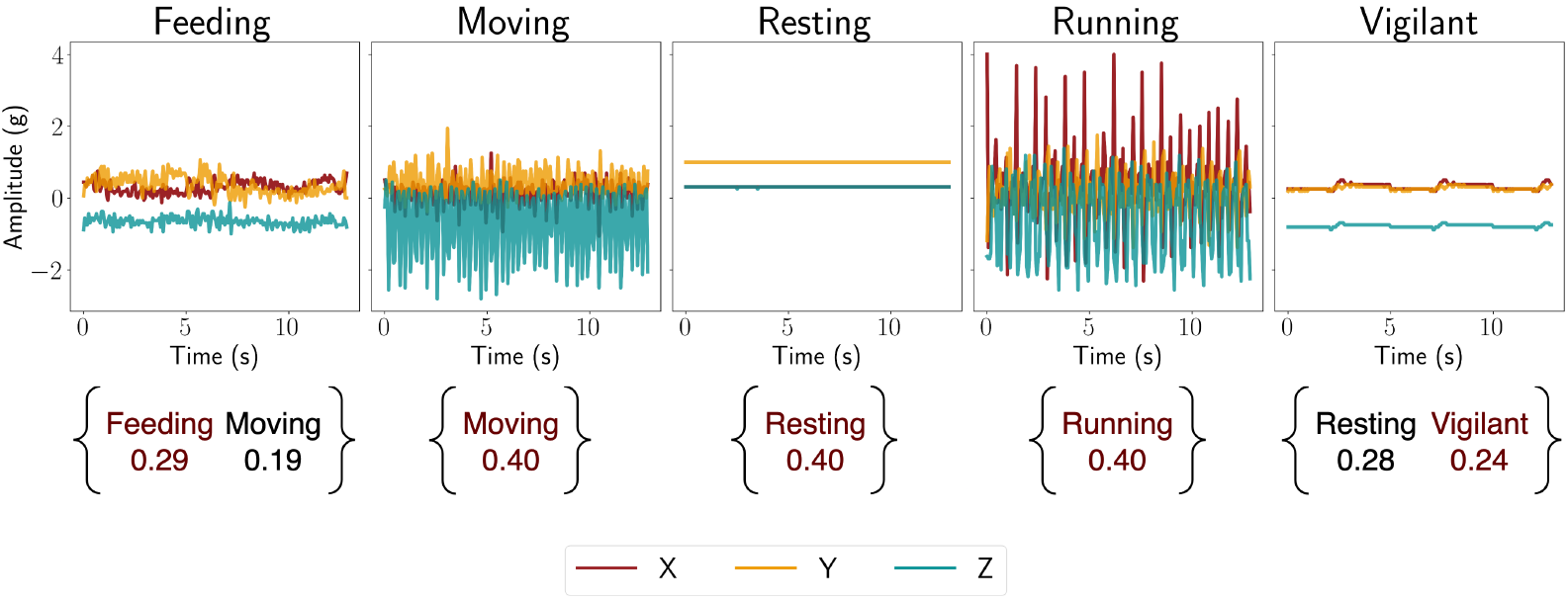
Regularised adaptive prediction sets (RAPS) of a random test sample from each behaviour class. RAPS refers to a prediction set which contains the true class with a probability of 1 ™*α* = 0.95. The numbers associated with each behaviour in the prediction set are the estimated and calibrated probability of success of that class. Here, the most likely behaviour in the prediction set is in blue and the remaining behaviours are in pink. Notice that the model estimates the true class with the highest probability for each behaviour.

Model training was carried out on a CUDA-enabled GPU machine with 12 GB of GPU memory, 64 GB of CPU RAM, and 24 virtual CPU cores. Creating the train-test split and fixed duration windows from the matched behaviour-acceleration pairs took approximately 2 minutes. This resulted in train, validation, and test set sizes of 14,978, 3,745, and 4,645, respectively. For a model configuration with 887,237 trainable parameters, the entire process of training took approximately 3 minutes. See Appendix F.2 for more details on computing.

After training ℳ_1_ on rebalanced training data, we fit the conformal model ℳ_2_ to the predictions of ℳ_1_ on the held-out validation set with a desired confidence level of 0.95. The overall performance of the prediction model ℳ_1_ and conformal model ℳ_2_ on various train-test splits is reported in the next section based on the evaluation metrics discussed in Section 2.4.2.

### 4.5 Testing Model Robustness to Distribution Shift

To test the model’s robustness against distribution shifts that may occur between training time and after implementation in the real world, we mimicked biologically realistic shifts using the metadata associated with our data. We created four experimental setups. Within the ‘No Split’ setup, we randomly divided our data into training and testing sets in an 80 − 20% ratio, maintaining class distributions across each set. Within the ‘Interdog’ setup, we trained our models with data from four dogs and reserved data from the fifth dog for testing. Within the ‘Interyear’ setup, we trained on data from one year (2021) and tested the data on a separate year (2022). Finally, within the ‘InterAMPM’ setup, we trained the model on data collected in the morning and tested the model on evening data. Additional distribution shift scenarios that could also be considered in similar systems could include, for example, assessing the impact of terrain characteristics (e.g., slope or vegetation type) on classification by splitting observations from distinct terrain types into separate train and test sets. Moving and running behaviours were broad categories that encompassed a range of movement speeds, from slow-paced walking to high-speed sprinting. As both categories included a range of speeds in the training and test data, model accuracy was unlikely to be strongly influenced by variation in movement speed within these categories, and they were not included within distribution shift analyses.

The gap between the train and test distributions was quantified using the sliced Wasserstein distance (see Appendix D.2). As shown in Table 3, the largest distribution shift occurred in the Interyear split, followed by Interdog and InterAMPM. For each experimental setup, the model was trained on the corresponding train data, and hyperparameter tuning was performed independently (see Appendix F).

**Table 3:** Dataset sizes, sliced Wasserstein distance between train and test data, and evaluation metrics (as described in Section 2.4.2) for the best-tuned classification ℳ_1_ and conformal ℳ_2_ models for the four distribution shift splits. For the metrics accuracy, precision, recall, and F1 score, the most likely class based on the scores output by ℳ_1_ are considered. This most likely class constitutes the Top-1 prediction set, and its accuracy is reported as Top-1 coverage. The RAPS coverage and RAPS average size assess the performance of ℳ_2_, which is fitted on the validation set. To compare model performance between the validation (same distribution as the training set) and testing sets, the metrics are presented side by side within the brackets.

| Evaluation Metric | No Split | Interdog | Interyear | InterAMPM |
| --- | --- | --- | --- | --- |
| Train set size | 14978 | 13104 | 9528 | 13712 |
| Validation set size | 3745 | 3277 | 2382 | 3429 |
| Test set size | 4645 | 6987 | 11458 | 6227 |
| Sliced Wasserstein Distance | 0.11 | 0.23 | 0.24 | 0.12 |
| Precision (val, test) | (0.89, 0.90) | (0.94, 0.84) | (0.94, 0.88) | (0.92, 0.90) |
| Recall (val, test) | (0.93, 0.94) | (0.95, 0.91) | (0.94, 0.91) | (0.92, 0.92) |
| F1 score (val, test) | (0.91, 0.92) | (0.94, 0.87) | (0.94, 0.89) | (0.92, 0.91) |
| Top-1 coverage (val, test) | (0.92, 0.93) | (0.93, 0.91) | (0.94, 0.90) | (0.94, 0.90) |
| RAPS coverage (val, test) | (0.94, 0.95) | (0.95, 0.93) | (0.95, 0.92) | (0.95, 0.92) |
| RAPS avg size (val, test) | (1.15, 1.15) | (1.10, 1.11) | (1.07, 1.10) | (1.06, 1.07) |

Table 3 presents model performance across all evaluation metrics described in Section 2.4.2 for the four experimental setups. As expected, performance generally declined with increasing distributional shift, with the largest drop from validation to test performance observed in the Interyear split; see Top-1 coverage (accuracy). The Top-1 coverage on the test data ranged from 0.90 for Interyear (representing the greatest distribution shift) to 0.93 for No Split (minimal shift). We note that the prediction sets notably improved the coverage of the true label, compared to the Top-1 prediction, across all experimental setups. For example, under No Split, coverage increased from 0.93 for the Top-1 prediction to 0.95 using RAPS. These results suggest that in settings with negligible distribution shift, RAPS effectively yields prediction sets that contain the true label for ≈ 95% of test observations. However, when the distribution shift is substantial, the added coverage provided by RAPS over Top-1 prediction is reduced.

### 4.6 Temporally Smoothed Classification

The final step of the pipeline, following model training and selection, involves applying the *trained* model to continuous streams of largely unlabeled accelerometer signals, segmented into fixed-duration windows. This is followed by smoothing the softmax scores produced by ℳ_1_. As described in Section 2.4.3, smoothing is conducted in two steps. First, the model is applied to consecutive windows of acceleration data, each 12.94 seconds long, along the temporal axis to generate a sequence of softmax scores. Second, this sequence of scores is smoothed by averaging the scores within an averaging window of length *s*. Thus, higher values of *s* represent greater degrees of smoothing and predictions on longer time windows, i.e., coarser predictions. The averaging window is then moved by *t* steps to obtain consecutive average scores. We used window sizes *s*, ranging from *s* = 25, suited for the detection of shorter-duration behaviours such as feeding, to s = 100, suited for longer-duration behaviours, such as resting (Figure 9).

**Figure 9.**
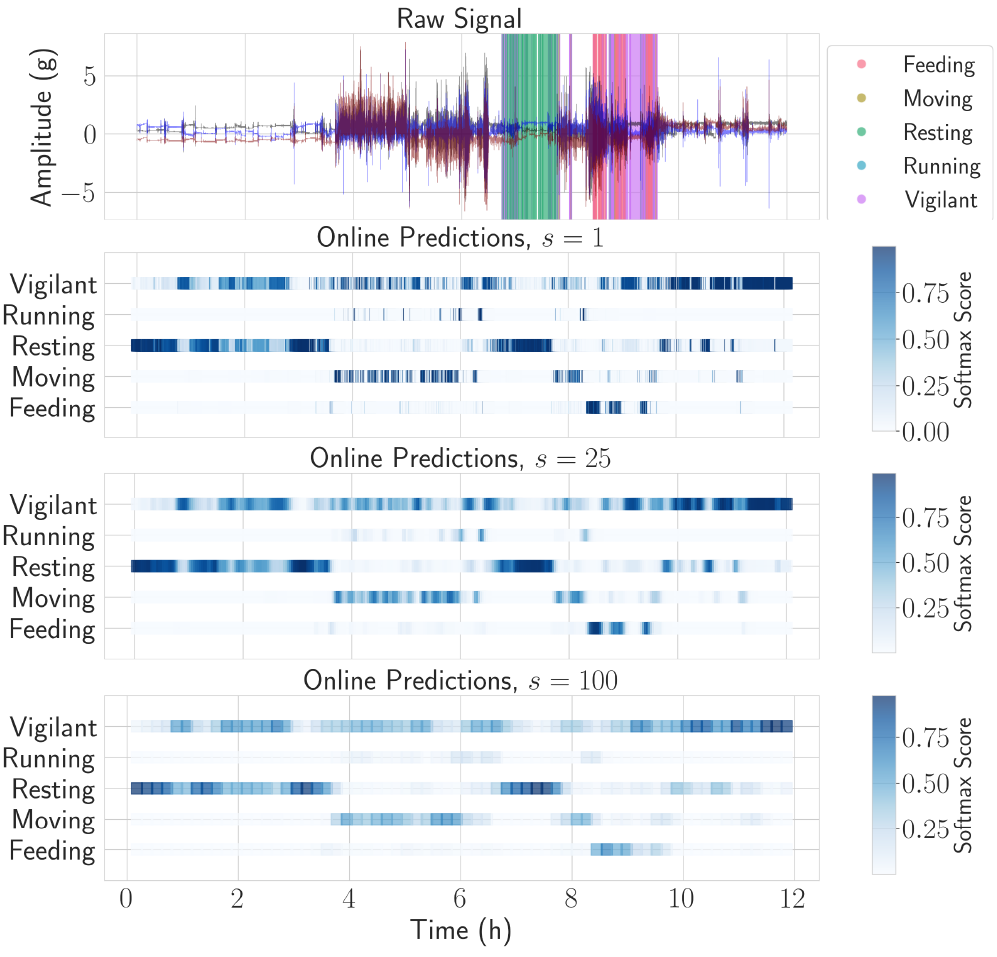
Smoothed classifications for a sample 12-hour African wild dog accelerometer segment. The top plot displays the original accelerometer signal, with shaded panels representing known labelled behaviours. The subsequent three plots illustrate the predicted probabilities of the different behavioural classes with no (*s* = 1), moderate (*s*=25), and high (*s*=100) average window lengths.

Ultimately, these temporally smoothed predictions could then be generated for entire accelerometer datasets, including months or years of data, enabling researchers to derive long-term behaviour timelines. For example, our pipeline could be applied to all half-day acceleration segments collected across deployment periods, producing continuous sequences of smoothed predictions. Researchers could then summarise behaviour frequency, timing, or transitions at biologically relevant scales (e.g. diel cycles, seasonal changes), supporting population-level ecological inference from fine-scale sensor data, even when direct observation is not possible.

## 5 Discussion

In this paper, we have introduced a novel approach for animal behaviour classification, applicable across taxa, that integrates machine learning with statistical inference to address common challenges and limitations of machine learning behaviour classification in ecology. Our approach allows users to evaluate and mitigate common issues in ecological datasets, including class imbalances and distributional shifts. Moreover, beyond providing single most-likely behaviour predictions, which typically lack probabilistic guarantees, our method offers robust sets of predicted behaviours that contain the true behaviour with a user-defined confidence level. Additionally, we incorporate a technique to smooth erratic behavioural predictions, which are often the result of fluctuating acceleration signals.

Our results highlight the utility of flexible class resampling techniques in improving animal behaviour classification accuracy, particularly in datasets with significant class imbalances. By combining undersampling and oversampling during model training, our framework enables researchers to achieve an optimal class distribution tailored to their study-specific needs, such as optimising precision or recall while minimising the biases introduced by excessive resampling Haixiang et al. (2017). For example, in our heavily imbalanced African wild dog dataset, rebalancing improved model recall by up to 29.2% when the class distribution was resampled to a mildly imbalanced state, with performance gains plateauing beyond this point. Accurate classifications for under-represented behaviours, such as feeding, more than doubled as a result. The severity of class imbalances and optimal resampling thresholds are likely to vary across populations and may be particularly severe for cryptic, wide-ranging, aerial, or marine species, where key behaviours can occur rarely or during periods when direct observation of individuals is impossible. For example, foraging behaviours represented only 0.49-1.46% of the total data for training seabird behaviour classifiers in Otsuka et al., 2024, whilst in arctic foxes, foraging (caching behaviours) represented 7.59% of collected data in Clermont et al., 2021. In cases where labelled samples are particularly sparse, resampling may yield little additional performance gains, and advanced data augmentation techniques may be required to improve model performance, such as artificially introducing noise into existing data or generating completely new data instances with AI (e.g., Luleci et al. (2022)). In such cases, researchers could expand upon our open-source pipeline and incorporate data augmentation techniques alongside resampling to optimise the extent of augmentation and resampling within a unified framework.

Quantifying and reporting uncertainty is a cornerstone of ecological research, and our pipeline is among the first to explicitly quantify uncertainty in animal behaviour classifications. By extending predictions from single behaviour classifications to prediction sets, we provide a framework analogous to the use of confidence intervals in statistical analyses rather than defaulting to binary outcomes of presence or absence of the behaviour of interest. Our approach thus provides a method to report and acknowledge model uncertainty within behavioural classification frameworks, which will become increasingly important as these methods become increasingly used within ecology and models are ported across different study systems. It is important to note that the conformal model is agnostic to the base classification model. While we used a 1D CNN architecture, conformal prediction can be applied to any base classification model that produces softmax scores (used to minimize the cross-entropy loss). In addition to providing a means to communicate uncertainty, which is important for independent evaluation and interpretation, our approach also provides a mechanism for researchers to propagate uncertainty from behaviour classifications into downstream analyses, such as quantifying how environmental (e.g., weather, human disturbance) or biological (e.g., social, physiological) processes affect animal behaviours. Moreover, due to training data needs and the questions of interest, in many cases, only a subset of species behaviours will be represented within training sets. In these cases, unrepresented behaviours will be classified into behaviours upon which the models were trained. The advantage of our prediction sets is that in such cases, assigned scores are likely to be closer across multiple categories, and prediction sets are likely to be larger, reflecting model uncertainty.

Uncertainty quantification additionally allows researchers to balance precision and confidence in behaviour classifications according to their study needs. As the user-chosen likelihood increases, so will the size of the prediction set and the confidence that it contains the true behaviour, and vice versa. As such, while higher likelihood thresholds will improve the likelihood of sets containing the true behaviour, they can also complicate downstream analyses by yielding a wider set of predicted behaviours. As a rule of thumb, we suggest likelihoods of 0.90 or 0.95 may be most familiar and suitable within ecology as they mirror the probability values widely used in null hypotheses testing across ecology (Castilho and Prado, 2021). We further recommend that researchers adopt the language of uncertainty when reporting model results, such that they indicate *moderate* or *strong* evidence for the prediction set containing the true behaviour based on 0.90 or 0.95 likelihood thresholds, respectively (see Muff et al., 2022). Ultimately, we see value and a need in developing shared standards around how prediction sets and their uncertainties are reported and used in ecological inference. Our pipeline provides one potential framework, but broader discussions, particularly on validation thresholds, reporting conventions, and ecological interpretation, will be important for developing shared standards that move the field forward.

Our pipeline also introduces a technique to evaluate model responses to distribution shifts, which can significantly impact model performance in real-world applications. Collecting labelled training data from animals in the wild can be logistically challenging and expensive. As a result, labelled ecological datasets often represent a narrow subset of the conditions in which the models will ultimately be applied. For example, labelled data collection may be carried out over relatively narrow time windows (Clermont et al., 2021), be constrained to an easily observable subset of the population (Chakravarty et al., 2019), or use captive animals (Rast et al., 2020). In our case study, we checked for potential sources of distributional shifts across years of data collection, study individuals, and diurnal periods; the lack of significant distributional shifts across these data subsets provides greater confidence in the validity of our models. While we provide tools for ecologists to assess model robustness to distributional shifts during model development, this study does not implement methods for mitigating performance decline under such shifts. This area of machine learning, known as domain adaptation, focuses on developing models that can generalise from a source data distribution to a different but related target distribution. Although several domain adaptation techniques for time series classification have been reviewed (Shi et al., 2022; Fawaz et al., 2023), their application within ecological research remains largely unexplored. Pending broader adoption of domain adaptation, we recommend a practical validation strategy for new datasets or deployments: utilising independently collected labelled video or audio segments. In such cases, we suggest comparing the observed behaviour from the audio or video against the model’s prediction set, rather than just the top-ranked prediction. For example, if a behaviour observed in a video is present in the model’s prediction set for that window, this constitutes a successful identification under the chosen confidence level. In cases where severe distribution shift is present, an appropriate domain adaptation technique or additional training data that captures the behavioural variability across the population of interest will likely be needed. Longitudinal monitoring is particularly important for long-term studies as plasticity in animal behaviours, updates in sensor technology, and the movement of new animal phenotypes into populations could lead to distributional shifts that undermine model performance.

We also provide a flexible approach to smoothing out noisy and anomalous behaviour classifications, drawing on techniques widely used in other fields such as signal processing and computer vision (Hamilton, 2020). Our smoothing technique allows users to filter out unlikely behavioural transitions by adding temporal context from surrounding classification windows into the classification process. For example, in our case study system, rapid and repeated transitions between feeding and running, or very short isolated feeding events, are biologically unlikely. Smoothing over a temporal window substantially reduced the frequency of abrupt and unlikely transitions between different predicted behaviour classes. The specific nuances of the study system and research questions of interest will dictate the intensity of smoothing required. Users should incorporate biological knowledge, such as typical behaviour durations, when selecting window lengths (analogous to smoothing intensity) to avoid filtering out key behaviours of interest. Larger window lengths will increase smoothing, reducing noise but risk potentially biasing against shorter behaviours. In contrast, shorter windows retain more noise but may inflate estimates for shorter duration behaviour categories. To help navigate this trade-off, we provide guidelines for selecting window lengths in the methods and for implementing it in our code. We also recommend conducting sensitivity analyses on window thresholds to evaluate and mitigate any biases that may arise from the smoothing.

To support the adoption of our pipeline, we provide open-source code, a step-by-step worked example, and recommendations for key decisions (e.g., parameter selection) that users will have to make. Whilst our method is implemented using tri-axial accelerometer data, this code could be added for bi-axial accelerometers. Recognising that hardware requirements may limit accessibility, we have developed our pipeline to function on a wide range of devices, from personal computers to GPU-equipped workstations. The largest impact on compute times comes from the availability of GPU support on machines. For example, model training (the most compute-intensive process) was 150 times quicker on GPU-equipped devices (see Appendix F.2 for more details). Ultimately, the compute time required to run our pipeline will vary across model parameters, data volumes, and computing hardware, but is likely to be on the order of minutes for studies with similar hardware and data volumes as our case study. For researchers without access to local hardware, the code can be executed seamlessly within a Google Colab notebook, enabling users to run the pipeline directly in their browsers.

## Conclusion

The use of accelerometers to classify animal behaviours is becoming increasingly widespread in animal ecology. Our study introduces a flexible, open-source approach that addresses common challenges, including class imbalances, distribution shifts, and uncertainty quantification. Using this approach, we demonstrated significantly improved predictions along with associated uncertainty metrics in African wild dog behaviour classification, particularly for rare and ecologically significant behaviours such as feeding, where correct classifications more than doubled following resampling. Looking forward, the integration of multiple sensor modalities, such as accelerometer, gyroscope, and GPS data, during model training offers intriguing opportunities for further improving model performance and expanding the range of detectable animal behaviours *in situ* (Castillo et al., 2014; Batpurev et al., 2021; Mao et al., 2021), and could be readily built into our pipeline. Our approach represents a key step towards advancing the burgeoning use of machine learning to remotely observe around-the-clock behaviours of free-ranging animals in their natural environments.

## Acknowledgements

We thank the Botswana Ministry of Environment, Wildlife, and Tourism for permission to conduct this research under research permit ‘ENT 8364 XLIX (38)’ to B.A. We thank the many researchers and collaborators associated with Botswana Predator Conservation for their invaluable contributions in the field and volunteers within the Abrahms lab for helping to annotate data. This work was funded by NSF DMS-2023166, CCF-2019844, DMS-2134012, NSF IOS (#2337405), the University of Washington Royalty Research Fund, eScience Institute, Washington Research Foundation, and the David and Lucile Packard Foundation.

## APPENDIX

[11pt]article [utf8]inputenc [left=1in,top=1in,right=1in,nohead,bottom=1in]geometry setspace [utf8]inputenc [T1]fontenc hyperref lineno url booktabs amssymb, amsthmbbm commath amsfonts amsmath nicefrac microtype graphicx subcaption multirow natbib doi lscape abstract [ruled,vlined]algorithm2e [dvipsnames]xcolor

Claim Lemma Fact Result Assumption Corollary Remark xr main

## SUPPLEMENTARY MATERIAL

### A Convolution Neural Networks

Here we described the architecture of a convolution layer in detail. Each convolution layer consists of three steps: 1D convolution, max pooling, and activation. The input to a 1D convolution step is of shape (*C*_in_, *T*), where *C*_in_ represents the number of input channels, and the output is of shape (*C*_out_, *T*), where *C*_out_ represents the number of output channels. For accelerometry data, the initial convolution layer starts with *C*_in_ = 3, so that we interpret each spatial axis as a channel. The number of output channels may not have such an interpretation, so it is thought of as a hyperparameter for the model’s architecture. Each output channel *i* (for *i* = 1, …, *C*_out_) is computed using the filter *W*_*i*_. The filter is a (*C*_in_, *h*) dimensional tensor where *h* is the kernel (or filter) size of the convolution. Let *y* represent the output from the 1D convolution, *y*_*i*_[*t*] denote the *t*-th element of the *i*th output channel, and *x*[*t*_1_ : *t*_2_] denotes the segment of *x* along the second axis starting from *t*_1_ and ending at *t*_2_. Then, *y*_*i*_[*t*] is obtained as

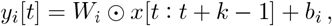

where ⊙ performs elementwise multiplication of two matrices, followed by the summation of their elements. Consequently, the output from such 1D convolution is of shape (*C*_out_, *T*).

The feature map highlights the presence and location of patterns or features detected by the filter. Choosing the right kernel size is essential, as it depends on factors like sampling frequency and duration of behaviours in the target species. The kernel size dictates the duration over which features are extracted. For species with slower sampling frequencies, larger kernels are more effective in capturing gradual changes in accelerometer readings. For example, to convolve over *u* seconds, the kernel size would be *uf*, where *f* is the sampling frequency. The kernel size is a hyperparameter that can be optimised by training the model on a range of values and selecting the one that yields the best performance on a validation set. The 1D convolution is followed by a max pooling layer. Max pooling extracts the maximum value within a specified window of a feature map. Using a window size of 2, we halve the size of the feature map along the temporal axis. The output is then passed through a rectified linear unit (ReLU) activation function, applying ReLU(*u*) = max(0, *u*) element-wise.

#### Model size considerations

The 1D CNN architecture is better suited than fully connected neural networks for extracting features from raw time series data from a memory point of view due to weight sharing (LeCun et al., 2015). The number of learnable parameters in each convolution layer of a 1D CNN depends on the number of CNN layers, input and output channels, and the kernel size, but is independent of the input signal dimension. This is beneficial when working with raw time series data, where input dimensions can be very large depending on the chosen window size for training. Conversely, in fully connected neural networks, the number of parameters grows with the input dimension.

### B Details for Methods

#### Creating fixed duration windows

We set the fixed window size to the *p*th percentile of behaviour durations across all observations in the dataset, where *p* is a user-specified parameter whose selection is verified through sensitivity analyses comparing different values of *p*. For observations exceeding this window duration, we extract multiple windows by splitting the signal into multiple parts. For those shorter than the window, we repeat the signal until it reaches the required duration, similar to circular padding in signal processing, where the signal is looped at the beginning and end to achieve the target length. Looping (as opposed to padding with silence) ensures that the model does not spuriously use the length of the annotation to identify behaviour and relies primarily on the content of the recording. For more details on padding techniques in signal processing, refer to Schoeters et al. (2020).

### C Regularized Adaptive Prediction Sets

Conformal prediction is a statistical framework that quantifies uncertainty in the predictions made by any base prediction model. At the heart of conformal prediction lies the coverage guarantee stated mathematically in the context of multi-class prediction in (3). This coverage guarantee is distribution-free in the sense that it does not depend on the data-generating distribution. The regularized adaptive prediction sets (RAPS), proposed by Angelopoulos et al. (2020), is a conformal prediction method lying in the broad range of *split* conformal prediction methods in the multi-class classification setting. Refer to the monograph by Angelopoulos et al. (2023) for a detailed review.

#### Calibrating softmax scores

First, the conformal model converts the softmax scores output by ℳ_1_ into calibrated scores. Several calibration techniques, such as Platt scaling (Platt et al., 1999) and isotonic regression (Zadrozny and Elkan, 2002), map softmax scores to calibrated class probabilities. We use Platt scaling in our method and describe it in greater detail here. For an observation (*x, y*), let (*ŷ*_1_, …, *ŷ*_*K*_) denote the softmax scores output by ℳ_1_. Calibrating the softmax scores using Platt scaling involves learning univariate parameters *A* and *b* such that the transformed scores

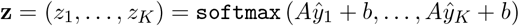

represent the calibrated probabilities. The parameters *A, b* are fit on a calibration dataset (disjoint from the training dataset used to train ℳ_1_) using logistic regression (Hosmer Jr et al., 2013). Refer to Platt et al. (1999) for more details on Platt scaling. While the calibrated softmax scores aim to provide better estimates of the conditional distribution Pr (*y* | *X)*, they offer no formal guarantees regarding model accuracy when predictions are made by selecting the label with the highest calibrated score. To address this, we proceed with the second step of the conformal model that provides such guarantees via prediction sets.

#### Constructing prediction sets

In general, conformal models aim to balance two goals while creating prediction sets: ensuring that the prediction set includes the true class with the desired coverage probability (satisfying (3)), and minimising the size of the prediction set to maintain concise predictions. Achieving this balance naturally introduces trade-offs, as larger sets naturally increase coverage but can become impractical. We use regularised adaptive prediction sets (RAPS) proposed by Angelopoulos et al. (2020), which balances these goals by incorporating a penalty for larger prediction sets while maintaining coverage guarantees (3). Concretely, given a calibration dataset 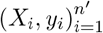, we proceed as follows.

1. We calculate the (calibrated or uncalibrated) softmax scores using the base classification model (here ℳ_1_). These softmax scores, denoted by (**z**_1_, …, **z**_*m*_) identify a heuristic notion of uncertainty over the set of classes.
2. Next, we define a conformal score function *E*(**z**, *y*) which takes the softmax scores **z** (of observation *X*) and a label *y* as input. A larger value of this score indicates worse alignment between the observation *X* (identified via **z**) and the label *y*.
3. Next, we compute the 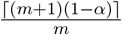 empirical quantile of conformal scores (**z**_1_, …, **z**_*m*_), denoted by 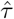.
4. Finally, we use this therehold 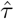 to form prediction sets for a new test observation (*X*^′^, *y*^′^) with softmax scores **z**^′^ as

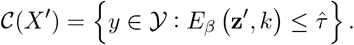

Different conformal prediction methods primarily differ in how they define the conformal score function *E*, which in turn governs their effectiveness. For RAPS, the conformal score function is randomized and parametrized using regularization parameters *β* = (*λ, k*_reg_). For an observation (*X, y*) with **z** denoting its sorted softmax scores (in descending order), let 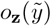 be the rank of the label 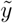 in **z** – e.g., if 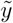 is the third most probable label, then 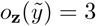. The conformal score is then defined as:

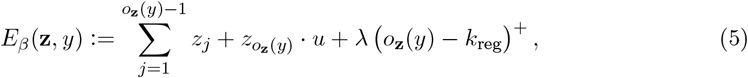

where *u* is a random sample from Unif(0, 1) and (*a*)^+^ denotes the positive part of *a*. The goal is to have larger values of conformal scores indicate worse alignment between *X* and *y*. Let us unwrap the score equation *E*_*β*_ in (5). The first term 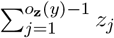 increases as *y* moves from most probable to least probable class according to the ordering in **z**. The second term 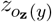 is a randomized term and handles the fact that that the value of the score can discretely jump with the inclusion of *y*, and the final term *λ* (*o*_**z**_(*y*) − *k*_reg_)^+^ is a regularization term that increases as the ranking of *y* increases, or equivalently, as *y* becomes more improbable according to the scores **z**.

Incorporating the score function *E*_*β*_ into our framework, Algorithm 1 details the computation of 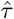, while Algorithm 2 describes how to construct RAPS using the fitted 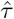 for a new test sample.

##### Algorithm 1

RAPS Conformal Calibration

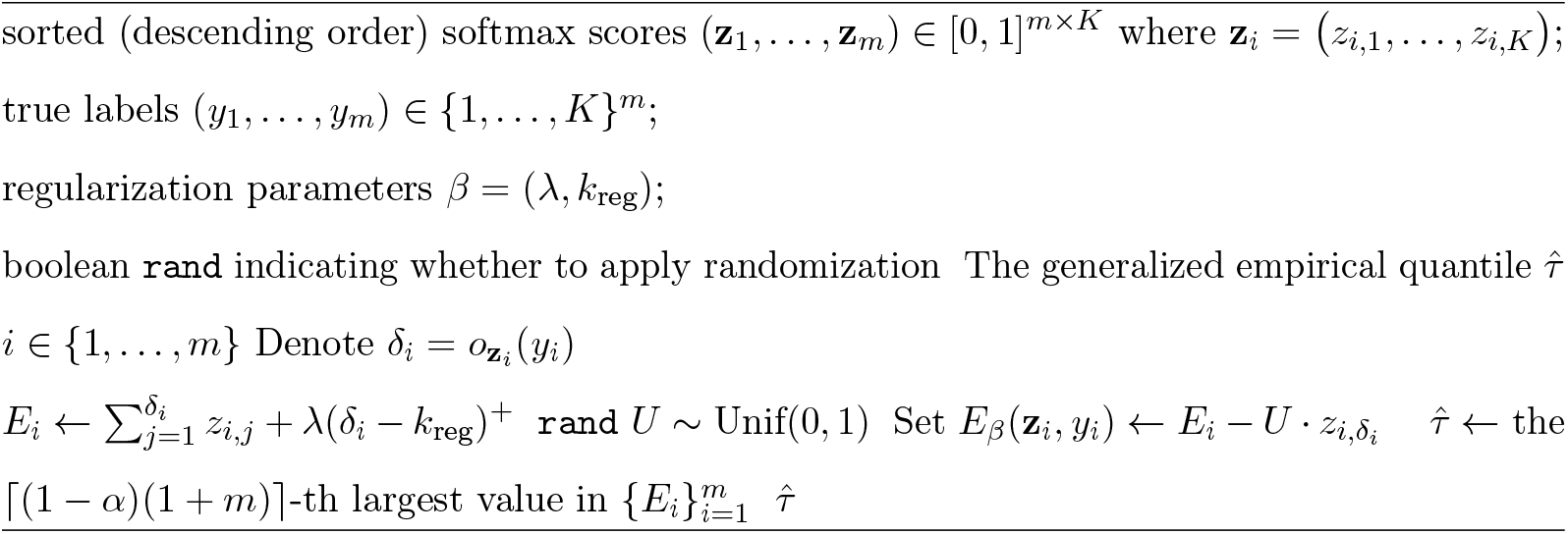

##### Algorithm 2

RAPS Prediction Sets

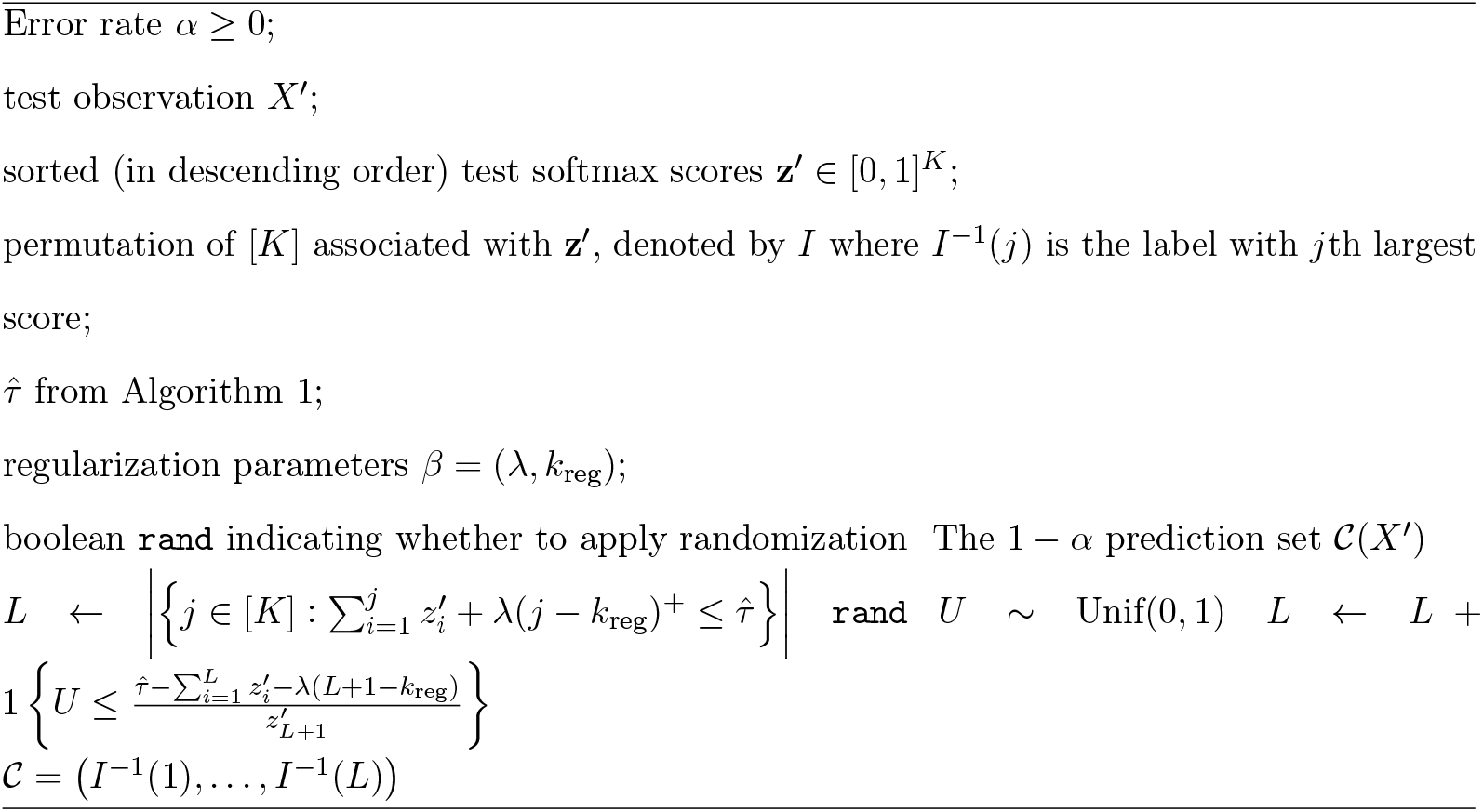

## D Simulation Details

### D.1 Hierarchical Model for Simulations

We use a hierarchical model, constructed as a mixture of sinusoids, to sample the acceleration data for each behaviour class and spatial axis. The model for acceleration signal *X*_*a*_(*t*), along axis *a* (out of *x, y, z* axes) given a behaviour label *y* is given by:

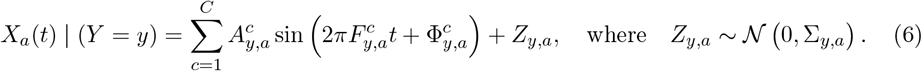

Thus, the signal is modeled as a sum of *C* (random) sinusoids with added Gaussian noise. For each behaviour *y* and axis *a*, the parameters of the *c*th sinusoid are the amplitude 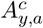, frequency 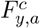, and phase shift 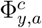. The parameter for the Gaussian noise is the variance Σ_*y,a*_. These parameters are randomly perturbed from fixed values as follows:

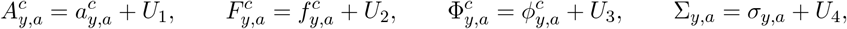

where *U*_*i*_, *i* ∈ {1, …, 4} are independent and identically distributed random variables. In summary, the parameters 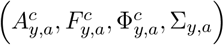 are defined as uniform perturbations (in the interval [−*τ, τ*]) of constants 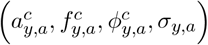. Throughout the simulations reported in the following subsections, we set *C* = 3 and *τ* = 0.35. The specific hyperparameter values of 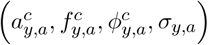 of (6) for the training set are detailed in Table 4.

**Table 4:**
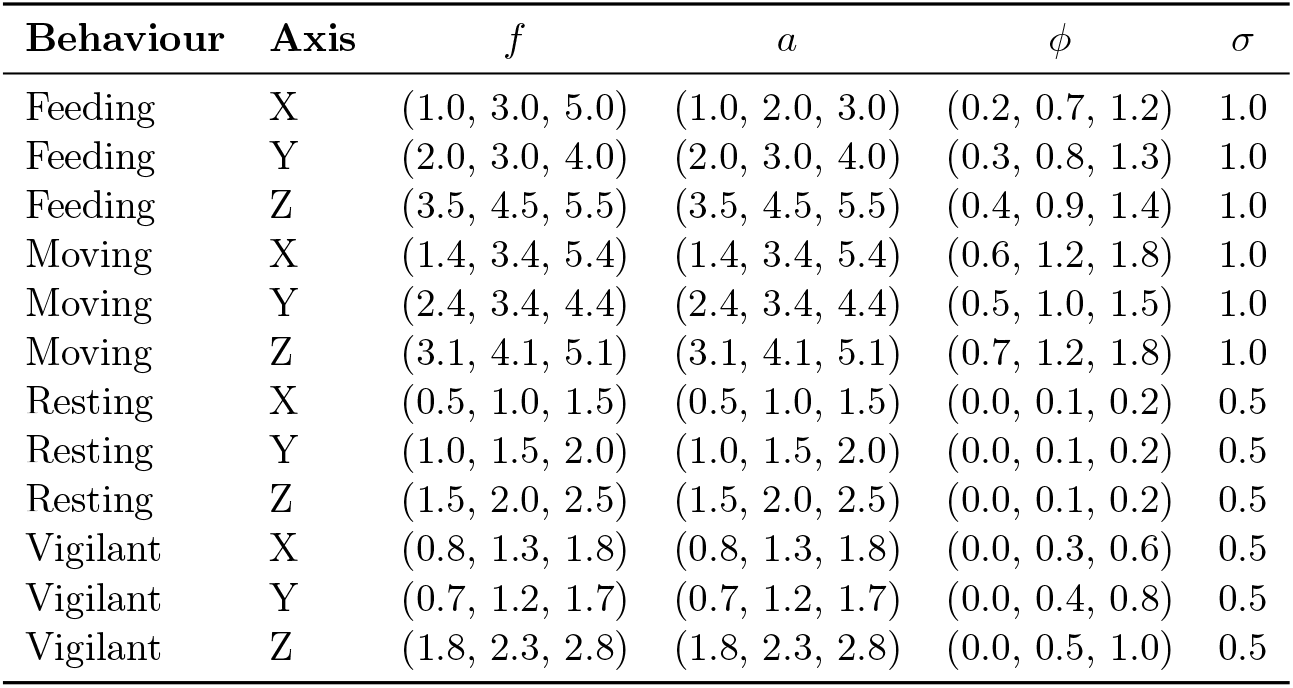
Simulation constants used to generate tri-axial acceleration signals for each behaviour and axis.

| Behaviour | Axis | $f$ | $a$ | $\phi$ | $\sigma$ |
| --- | --- | --- | --- | --- | --- |
| Feeding | X | (1.0, 3.0, 5.0) | (1.0, 2.0, 3.0) | (0.2, 0.7, 1.2) | 1.0 |
| Feeding | Y | (2.0, 3.0, 4.0) | (2.0, 3.0, 4.0) | (0.3, 0.8, 1.3) | 1.0 |
| Feeding | Z | (3.5, 4.5, 5.5) | (3.5, 4.5, 5.5) | (0.4, 0.9, 1.4) | 1.0 |
| Moving | X | (1.4, 3.4, 5.4) | (1.4, 3.4, 5.4) | (0.6, 1.2, 1.8) | 1.0 |
| Moving | Y | (2.4, 3.4, 4.4) | (2.4, 3.4, 4.4) | (0.5, 1.0, 1.5) | 1.0 |
| Moving | Z | (3.1, 4.1, 5.1) | (3.1, 4.1, 5.1) | (0.7, 1.2, 1.8) | 1.0 |
| Resting | X | (0.5, 1.0, 1.5) | (0.5, 1.0, 1.5) | (0.0, 0.1, 0.2) | 0.5 |
| Resting | Y | (1.0, 1.5, 2.0) | (1.0, 1.5, 2.0) | (0.0, 0.1, 0.2) | 0.5 |
| Resting | Z | (1.5, 2.0, 2.5) | (1.5, 2.0, 2.5) | (0.0, 0.1, 0.2) | 0.5 |
| Vigilant | X | (0.8, 1.3, 1.8) | (0.8, 1.3, 1.8) | (0.0, 0.3, 0.6) | 0.5 |
| Vigilant | Y | (0.7, 1.2, 1.7) | (0.7, 1.2, 1.7) | (0.0, 0.4, 0.8) | 0.5 |
| Vigilant | Z | (1.8, 2.3, 2.8) | (1.8, 2.3, 2.8) | (0.0, 0.5, 1.0) | 0.5 |

Each random observation is denoted by (*X, Y*) where *X* is the fixed duration tri-axial raw acceleration signal and *Y* is the associated behaviour label. For a fixed window length *T*, we have that *X* ∈ ℝ^3×*T*^ and *Y* ∈ {Feeding, Moving, Resting, Vigilant}. We assume that the sampling frequency is 16 Hertz and the fixed window duration is 30 seconds, and therefore *T* = 480. Then, for a given behaviour label *Y* = *y*, the acceleration signal is sampled according to the model (6).

#### D.2 Discussion on Distribution Shift Results

##### Simulation Setup for Distribution Shift Results

We choose the same class distribution (distribution of *Y*) for the test set and the distribution of *X* given *Y* again follows the model in (6) with modified model parameters, denoted by 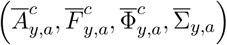, and distributed as

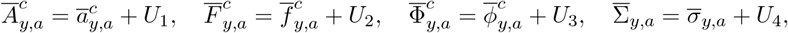

where again *U*_*i*_ are independent and identical distributed random variables with *U*_*i*_ ∼ Unif(−*τ, τ*) for *i* ∈ {1, …, 4}. Here, 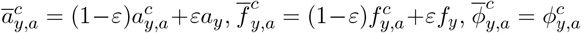 and 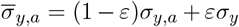. Thus, 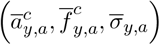 interpolate between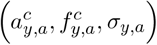 and *(a*_*y*_, *f*_*y*_, *σ*_*y)*_. The parameters (*a*_*y*_, *f*_*y*_, *ϕ*_*y)*_ are shared across behaviour pairs: feeding-moving and resting-vigilant and provided explicitly in Table 5.

**Table 5:**
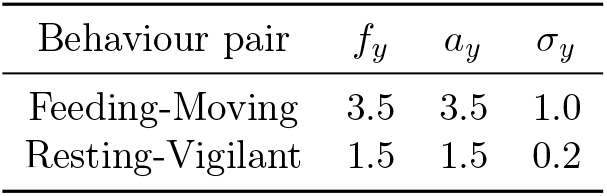
Simulation constants for test distribution with *ε* = 1 for the distribution shift simulation setup.

| Behaviour pair | $f_y$ | $a_y$ | $\sigma_y$ |
| --- | --- | --- | --- |
| Feeding-Moving | 3.5 | 3.5 | 1.0 |
| Resting-Vigilant | 1.5 | 1.5 | 0.2 |

##### Summary Statistics of Accelerometry Data

Following Nathan et al. (2012), each acceleration window of shape (3, *T*), where *T* is the window length, was summarized using 34 features. First, in addition to the raw acceleration along the *X, Y*, and *Z* axes, we defined a fourth axis representing overall dynamic body acceleration:

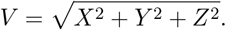

For each of the four axes (*X, Y, Z*, and *V*), we computed seven statistical summaries: mean, standard deviation, skewness, kurtosis, maximum, minimum, and lag-1 autocorrelation, yielding 28 features. We then computed the three pairwise Pearson correlations among the *X, Y*, and *Z* axes. Additionally, overall dynamic body acceleration (ODBA) was calculated and included as a summary statistic (Shepard et al., 2008; Wilson et al., 2006). Finally, we computed inclination *ϕ* = cos^−1^(*Z/V*) and azimuth *ψ* = tan^−1^(*Y/X*), and included their circular variances as features, resulting in a total of 34 summary statistics.

##### Sliced Wasserstein Distance

The *Sliced Wasserstein Distance* is a computationally efficient way to measure the dissimilarity between two probability distributions in high-dimensional spaces. Given two distributions *P* and *Q* on ℝ^*d*^, the idea is to project them onto one-dimensional subspaces defined by directions *θ* ∈ 𝕊^*d*−1^, where 𝕊^*d*−1^ denotes the unit sphere in ℝ^*d*^ (i.e., the set of all unit vectors *θ* ∈ ℝ^*d*^ such that ‖*θ*‖ = 1). For each direction *θ*, the one-dimensional Wasserstein distance *W*_2_(*P*_*θ*_, *Q*_*θ*_) is computed between the projected distributions *P*_*θ*_ and *Q*_*θ*_. The one-dimensional Wasserstein distance can be computed in closed form Villani (2021). The sliced Wasserstein distance is obtained by averaging these over all directions:

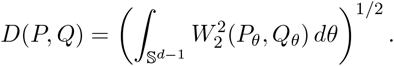

In practice, when only finite samples {*x*_1_, …, *x*_*n*_} from *P* and {*z*_1_, …, *z*_*n*_} from *Q* are available, aliced Wasserstein distance can be approximated by randomly sampling *K* directions *θ*_1_, …, *θ*_*K*_ ∈ 𝕊^*d*−1^, projecting the samples, sorting the resulting 1D projections, and computing empirical Wasserstein distances for each direction. The final estimate is then:

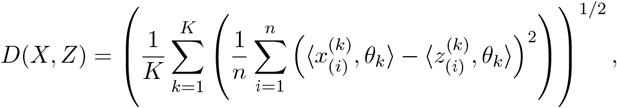

where 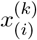 and 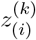 denote the sorted projections of the samples along direction *θ*_*k*_. This formulation enables efficient and scalable comparison of high-dimensional empirical distributions.

### D.3 Simulating 24 Hours of Acceleration-Behaviour Signal

To evaluate the prediction and conformal models on long-duration continuous data, we simulate 24 hours of acceleration signals representing different behaviours. Behaviour transitions are governed by a Markov kernel (Table 7), where the next behaviour is sampled from a distribution conditioned on the current one. Each row of the table specifies the transition probabilities from a given behaviour to all others. Each behaviour has a predefined average duration informed by ecological knowledge. The acceleration signals are generated using the hyperparameters listed in Table 6. A sample simulated day is shown in Figure 10, and is used in Sections 3.2 and 3.4.

**Table 6:** Transition kernel and average duration for simulating long durations of behaviour data.

| Behaviour | Transition Kernel |  |  |  | Average Duration (hrs) |
| --- | --- | --- | --- | --- | --- |
|  | Feeding | Moving | Resting | Vigilant |  |
| Feeding | 0.0 | 0.7 | 0.0 | 0.3 | 1 |
| Moving | 0.5 | 0.0 | 0.3 | 0.2 | 4 |
| Resting | 0.0 | 0.7 | 0.0 | 0.3 | 6 |
| Vigilant | 0.0 | 0.5 | 0.5 | 0.0 | 2 |

**Table 7:** Standard deviation of softmax scores with and without smoothing.

| Behaviour | No smoothening, $s = 1$ | | | | Smoothening, $s = 20$ | | | |
| --- | --- | --- | --- | --- | --- | --- | --- | --- |
|  | Feeding | Moving | Resting | Vigilant | Feeding | Moving | Resting | Vigilant |
| Feeding | 0.2085 | 0.1701 | 0.0078 | 0.1174 | 0.0709 | 0.0672 | 0.0132 | 0.0165 |
| Moving | 0.0041 | 0.1164 | 0.0428 | 0.1720 | 0.0179 | 0.0340 | 0.0219 | 0.0209 |
| Resting | 0.0009 | 0.1868 | 0.2138 | 0.1554 | 0.0001 | 0.0285 | 0.0367 | 0.0191 |
| Vigilant | 0.0072 | 0.1502 | 0.2864 | 0.2843 | 0.0235 | 0.0134 | 0.0252 | 0.0170 |

**Table 8:**
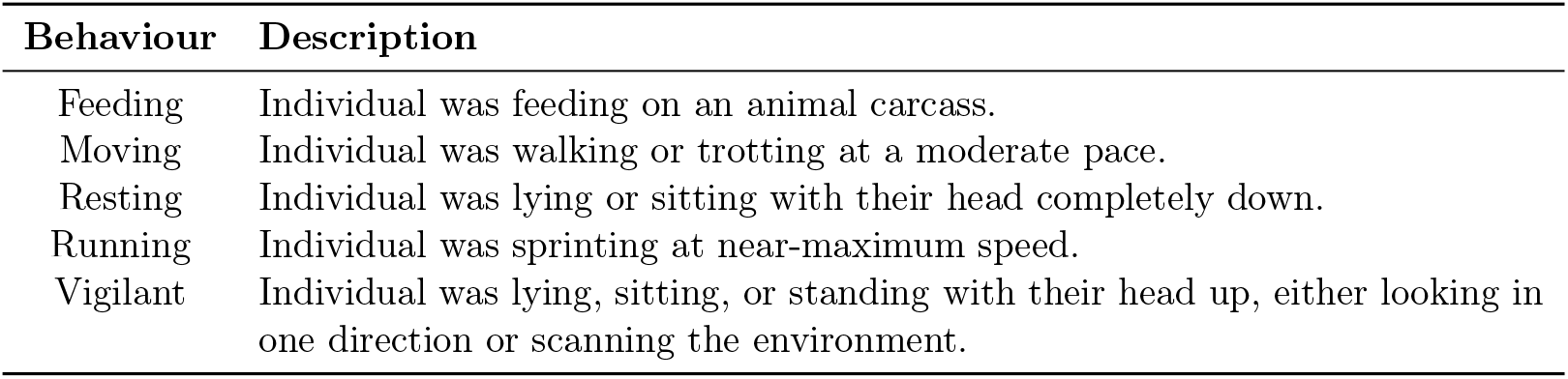
Individual-wise summary of labelled acceleration data including start and end date of matched data, number of unique days, duration of matched data, and total number of extracted windows of duration *w* seconds.

**Table 9:**
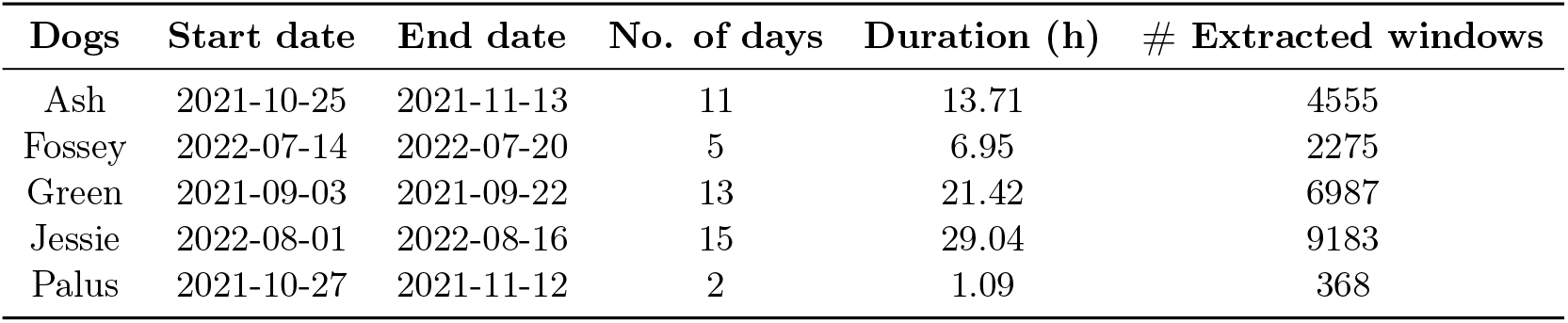
Individual-wise summary of labeled acceleration data including start and end date of matched data, number of unique days, duration of matched data, and total number of extracted windows of duration *w* seconds.

| Dogs | Start date | End date | No. of days | Duration (h) | # Extracted windows |
| --- | --- | --- | --- | --- | --- |
| Ash | 2021-10-25 | 2021-11-13 | 11 | 13.71 | 4555 |
| Fossey | 2022-07-14 | 2022-07-20 | 5 | 6.95 | 2275 |
| Green | 2021-09-03 | 2021-09-22 | 13 | 21.42 | 6987 |
| Jessie | 2022-08-01 | 2022-08-16 | 15 | 29.04 | 9183 |
| Palus | 2021-10-27 | 2021-11-12 | 2 | 1.09 | 368 |

**Figure 10.**
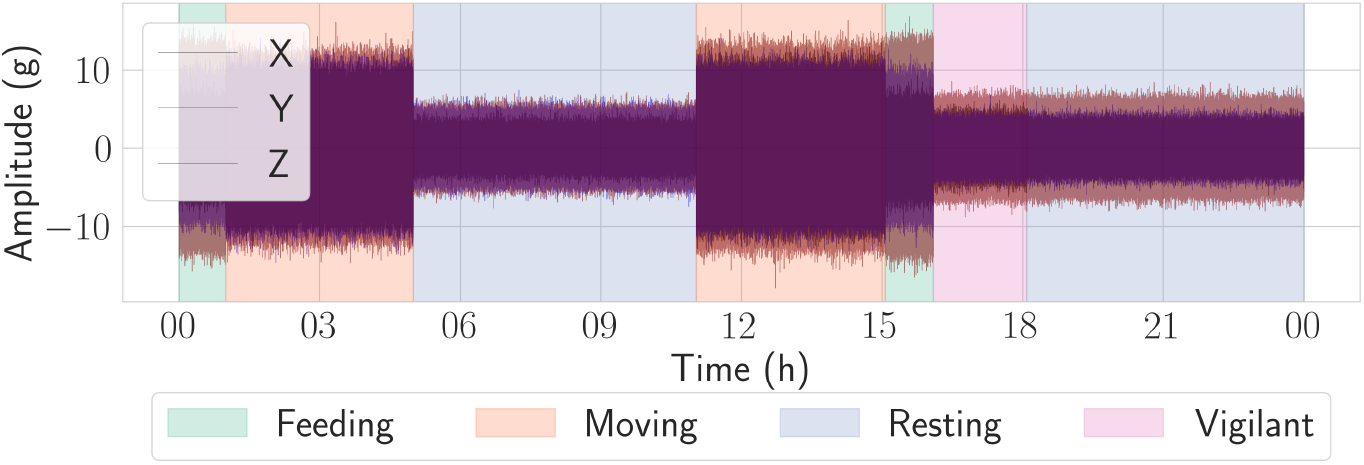
Simulated acceleration-behaviour signal for 24 hours duration

**Figure 11.**
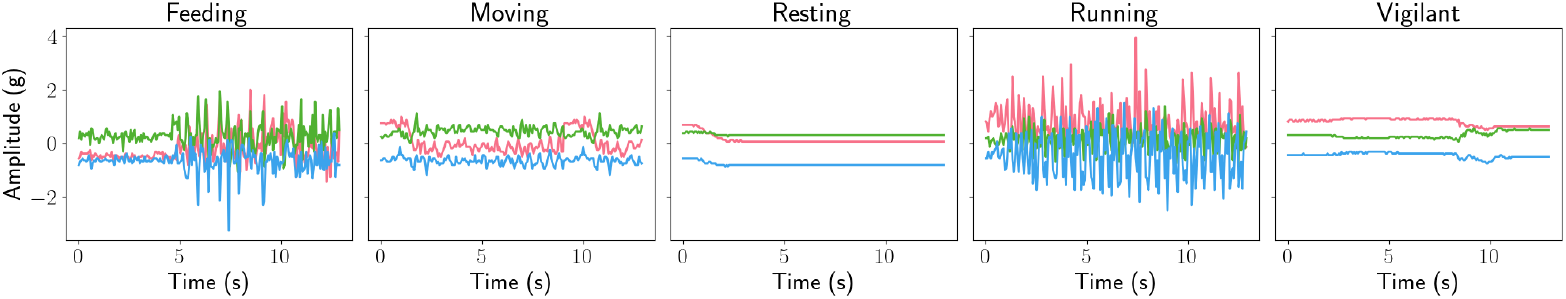
Accelerometry data trace plot for a randomly selected observation of each behaviour class, with a consistent window duration of *w* = 12.94 seconds.

### D.4 Discussion on Temporally Smoothed Classification Results

Here we discuss a quantitative way of assessing temporal smoothening via standard deviation of softmax scores for each true behaviour class. As discussed in Section 3.4, we expect softmax scores to be more stable (i.e. less variable) for behaviours the model is confident about. Figure 3 shows that the model frequently confuses the minority behaviours feeding and vigilant with the majority behaviours moving and resting, respectively. In particular, resting and vigilant are often confused with each other (Figure 6). As a result, we expect high score variability for behaviours the model struggles to distinguish, and low variability for behaviours it consistently predicts with low probability.

Table 7 reports the standard deviation of softmax scores across all predicted behaviours, grouped by the true class. In the left panel (no smoothing), moving exhibits high variability, regardless of confusion, likely due to corrupted signals being misclassified as moving. Applying temporal smoothing with *s* = 20 reduces this spurious variability, removing sudden peaks in moving scores, and yields more stable predictions, albeit over a longer temporal window (10 minutes). The remaining variability in the smoothed scores reflects genuine model uncertainty between similar behaviours, and can thus be used to construct more meaningful prediction sets.

## E Details on African Wild Dogs Data

### E.2 Details on African wild dog data collection process

We paired accelerometer data with 103.77 hours of time-stamped video collected from a handheld video camera recording from within an observation vehicle. African wild dogs within the study system were habituated to vehicles due to research and tourism activity within the area, and as such, no obvious changes in behaviour were observed due to the presence of vehicles. All collars had biodegradable cotton links (three layers of cotton), which degraded over time following environmental exposure, leading to remote collar detachment once completely degraded, or were manually removed and replaced with new collars following battery expiry. Accelerometer and audio data were manually downloaded after collecting tracking collars or detached audio recorders, respectively. We attached collars in collaboration with a Botswana-registered veterinarian who sedated animals using established immobilisation protocols (for more details, see Hubel et al. (2016a,b)). During collar attachment on sedated animals, we used vehicles to visually separate immobilised animals from other African wild dogs in the pack to minimise stress for the remainder of the group, which was typically resting less than 200 m away. After immobilisation, collared animals returned to the rest of the pack within 15 minutes. Our work received ethical approval from the University of Washington (IACUC #4514-01) and Botswana’s Department of Wildlife and National Parks (Permit # ENT 8364 XLIX (38)). No ill effects of collar deployments were observed.

#### Behaviour Details

Table 8 provides a description of the five behaviours chosen for classification. Table 9 provides an individual-wise summary of the 72.21 hours of labelled acceleration data used for model training.

## F Training Details

### F.2 Hyperparameter Finetuning

Figure 12 shows the precision, recall, and F1 score on the validation set, averaged across six seeds, for different hyperparameter configurations under all four experimental setups. Across experiments, the highest performance is consistently achieved by the most expressive model—5 convolutional layers, 64 output channels in the first layer, and kernel size 5. However, for the No Split experiment, increasing the number of convolutional layers from 3 to 5 yields only a marginal improvement. To balance performance and computational efficiency, we therefore adopt a CNN with three convolutional layers for the No-Split experiment.

**Figure 12.**
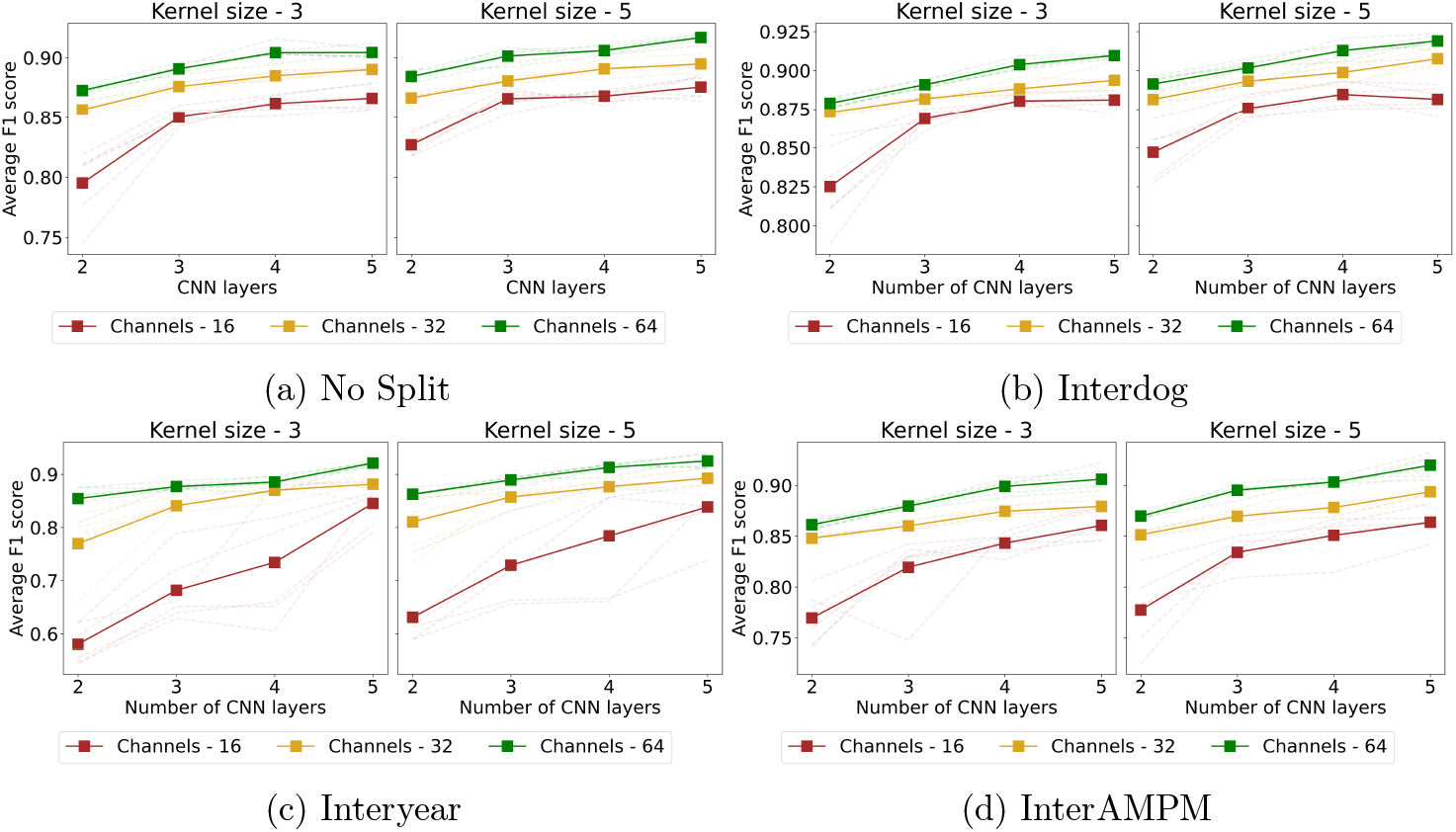
Hyperparameter tuning for the four experiments. Tunable parameters are the number of CNN layers, the number of output channels in the first layer, and the kernel size.

**Figure 13.**
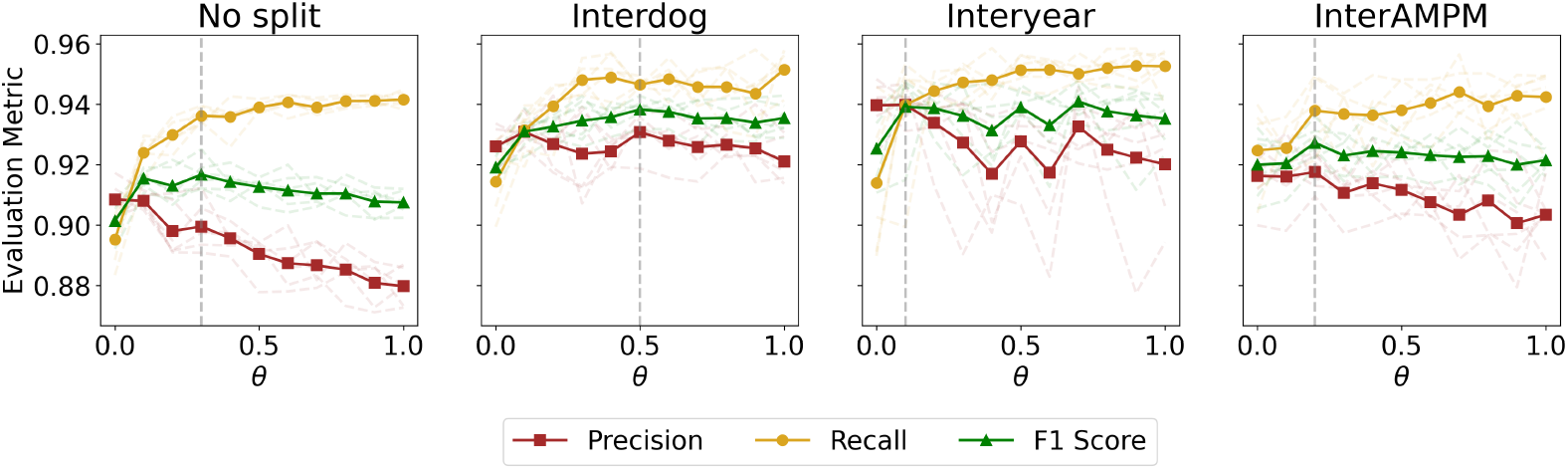
Tuning for rebalancing parameter *θ* for four experiments by comparing the precision, recall, and F1 score on the validation set. The number of output channels is fixed at 64, kernel size at 5, and the number of CNN layers is determined by optimal values from Figure 12. The metrics are averaged over six different training runs with different seeds.

### F.2 Compute Details

All experiments for the African wild dog case study were conducted on a GPU-equipped machine with CUDA support, featuring 12 GB of GPU memory, 64 GB of CPU RAM, and 24 virtual CPU cores. To assess the performance of our pipeline on machines without GPU support, we also ran the pipeline on a workstation with 32 virtual CPU cores and 125 GB of RAM. Additionally, the pipeline was tested on a personal Apple MacBook with an 8-core CPU and 16 GB of RAM. The time-consuming steps of the pipeline can be divided into three main processes. The first step, metadata creation, involves reading all half-day CSV files and extracting information such as year, individual identification, and the average temperature for each half-day. This step is CPU-bound and takes approximately 148 seconds to process a total of 4,276 half-day CSV files. The second step, creating training and testing splits using user-defined filters, takes around 70 seconds across all machines. The third step, creating fixed-duration windows from the filtered data, consistently takes about 25 seconds across all platforms. GPU support proves to be particularly advantageous during the final step - model training, where efficient parallelism for tensor computations accelerates the process. The machine learning model is built in the PyTorch framework. The model with 5 CNN layers, 32 output channels, and kernel size 5 has 887,237 trainable parameters. For a training set size of 14,978 samples and this model configuration with 887,237 trainable parameters, one epoch of training on the GPU-equipped machine takes an average of 2 seconds. In comparison, training on the workstation takes approximately 25 seconds per epoch, while the personal MacBook requires around 330 seconds per epoch.

